# Glycine receptor subunit-ß -deficiency in a mouse model of spasticity results in attenuated physical performance, growth and muscle strength

**DOI:** 10.1101/2020.07.15.180976

**Authors:** Cintia Rivares, Alban Vignaud, Wendy Noort, Bastijn Koopmans, Maarten Loos, Mikhail Kalinichev, Richard T Jaspers

**Affiliations:** Laboratory for Myology, Department of Human Movement Sciences, Faculty of Behavioral and Movement Sciences, Vrije Universiteit Amsterdam, Amsterdam Movement Sciences, Amsterdam, The Netherlands; Ipsen Innovation, Les Ulis, France; Sylics (Synaptologics BV), Amsterdam, The Netherlands

**Keywords:** spastic cerebral palsy, hereditary spastic paraplegia, triceps surae muscles, hyper-resistance, Glrb^spa^, m. gastrocnemius, m. soleus, ankle plantar flexor muscles, gait, motor function, PhenoTyper, Catwalk

## Abstract

Spasticity is the most common neurological disorder associated with increased muscle contraction causing impaired movement and gait. The aim of this study was to characterize physical performance and skeletal muscle function and phenotype of mice with a hereditary spastic mutation (B6.Cg-Glrb^spa^/J). Motor function, gait and physical activity of juvenile and adult spastic mice and the morphological, histological and mechanical characteristics of their soleus (SO) and gastrocnemius medialis (GM) muscles were compared with their wild-type (WT) littermates. Spastic mice showed attenuated growth, impaired motor function and low physical activity. Gait of spastic mice was characterized by a typical hopping pattern. Spastic mice showed lower muscle forces, which were related to the smaller physiological cross-sectional area of spastic muscles. The muscle-tendon complex length-force relationship of adult GM was shifted towards shorter lengths, which was explained by attenuated longitudinal tibia growth. Spastic GM was more fatigue resistant than WT GM. This was largely explained by a higher mitochondrial content in muscle fibers and relatively higher percentage of slow type muscle fibers. Muscles of juvenile spastic mice showed similar differences compared to WT juvenile mice, but these were less pronounced than between adult mice. This study shows that in spastic mice, disturbed motor function and gait is likely the result hyperactivity of skeletal muscle and impaired skeletal muscle growth, which progress with age.

## Introduction

Spastic paresis, due to acquired perinatal upper motor neuron lesion (i.e. cerebral palsy, CP) or genetic mutations (i.e. hereditary spastic paraplegia, HSP) accounts for the most common cause of movement disorders in children (Paneth et al. 2006). Children with spasticity experience over-activity of the stretch reflex, provoking resistance against extension of joints (hyper-resistance). CP occurs in about 2 to 3 per 1.000 liveborn children and 80% of these children suffer from spasticity (Himmelmann and Uvebrant 2018; Christensen et al. 2014; Cans 2000). Even though HSP is less common than CP, its prevalence is still about 3 to 10 per 10.000 liveborn children in Europe (Fink 2013; Mc Monagle, Webb, and Hutchinson 2002). Spasticity is most prominent in ankle plantar flexor and knee flexor muscles causing joint hyper-resistance against extension of these joints and a deteriorated gait. Typical ‘toe walking’ (also called ‘equinus’) and crouch gait patterns, as well as abnormal knee flexion during terminal swing phase are results of hyper-resistance (Goldstein and Harper 2001; Rosenbaum et al. 2006; Haberfehlner et al. 2018; Cooney et al. 2006; Rodda and Graham 2001).

During locomotion, excessive antagonistic co-contraction of spastic muscles, contribute to the net torque over target joints, requiring higher agonist muscle forces to execute particular movements (Damiano et al. 2000; Van Den Hecke et al. 2007; Unnitahn et al. 1996; Ikeda et al. 1998), which consequently increases the energy demand. Mobility, as well as social participation of children with spasticity are reduced (Bjornson et al. 2007; Zwier et al. 2010; Schenker et al. 2005; Sahoo et al. 2017), which ultimately implicates a lower daily physical activity of children with spasticity compared to their typical developing peers (Zwier et al. 2010; Schenker et al. 2005).

During development, the spasticity related hyper-resistance also may affect skeletal muscle growth and adaptation, as well as that of connective tissue and bony structures (Graham and Selber 2003; Huijing et al. 2013; Mathewson et al. 2015). Several studies have shown that longitudinal fascicle growth is impaired in children with spasticity, particularly in muscles of the lower limb (Mohagheghi et al. 2008; Wren et al. 2010; Gao et al. 2011; Barber et al. 2012). The underlying mechanism for this may be an impaired longitudinal muscle fiber growth as has been shown during development in children with SP (Barber et al. 2012; Haberfehlner et al. 2016), which indicates that serial sarcomere number must be likely lower in these fascicles. As a consequence, spastic muscles operate at higher sarcomere lengths than those of typical developed muscles, which may explain the increased passive resistance (Mathewson et al. 2015; Smith et al. 2011). Note that, since human muscles from the lower extremities have mainly a pennate morphology (Ward et al. 2009), longitudinal muscle growth also occurs by an increase in muscle fiber cross-sectional area (Bénard et al. 2011; Heslinga and Huijing 1990; Weide et al. 2015, Swatland, 1980). In addition to impaired muscle growth, structural alterations of collagen and extracellular matrix may contribute to muscle hyper-resistance, because of their incrementing effect on stiffness of muscle fiber bundles (Smith et al. 2011; Mathewson et al. 2014). However, whether the connective tissue content is increased in spastic muscle is subject to controversy and seems to differ between muscles (cf. De Bruin et al. 2014; Mathewson et al. 2015; Smith et al. 2011).

Clinical interventions aim to reduce muscle hyperactivity and increase the range of joint motion. Although, interventions are temporally successful, large variations in treatment outcomes have been reported (Bottos et al. 2003; Moore et al. 2008). Moreover, long-term functional limitations are recurrent (Fry et al. 2007). Improvement of interventions and development of new drugs require insight into mechanisms underlying spasticity. In vivo studies with typical developing children and children with spasticity are limited with respect to the level of detail of the underlying mechanistic causes.

Animal models of spasticity allow obtaining a more detailed insight into the multitude of determinants of hyper-resistance and impaired mobility. There are two types of animal models for spasticity, one based on acute trauma (e.g. transection, hemisection or injury of the spinal cord) and another on genetic mutations. In particular the spastic mouse model, mice with inherited spasticity by a spontaneous mutation, is a valuable model that presents symptoms of spasticity at an early age (2 weeks of life) and can reach an adult age (Chai 1961; Mülhardt et al. 1994). This makes the spastic mouse model suitable for investigation of the mechanisms underlying spasticity associated impairments in gait as well as testing of novel interventions to improve physical performance. However, the extent of behavioural changes and muscle function of this mouse model have not been quantified in detail.

Mice homozygous for the spontaneous spastic mutation (spastic; Spa) have an intronic insertion of a LINE 1 element in the inhibitory Glycine receptor ß-subunit gene (*Glrb*^*spa*^) which causes exon skipping and leads to a nearly complete down regulation of the expression of the ß-subunit (Mülhardt et al. 1994). Reduced glycine receptor inhibition gives rise to a different motor neuron firing pattern that is associated with overactivation of the muscle, caused by lack of fast inhibitory transmission due to lower Chloride content within the cells. The glycine receptor mutation in mice has been described to cause startle response, muscle rigidity, tremor and impaired righting, as well as impaired gait (Chai 1961), but these phenotypes have not been quantified in detail. Mutations in glycine receptor are also known to occur in humans, which causes hyperekplexia, causing exaggerated startle responses and increase muscle tone (Langosch et al. 1994).

The aim of this study was twofold:1) To quantitatively characterize gait, motor function and physical activity of spastic mice across post-natal development, at the age of 4, 6, 10 and 14 weeks and 2) To obtain insight in the mechanisms underlying spasticity associated limitations in physical performance. In particular, we assessed contractile force characteristics, and morphological and histological phenotype of two lower limb muscles (i.e. gastrocnemius, GM and soleus, SO) in juveniles (6 weeks) and adults (18 week). In addition, we assessed the expression of growth factors in the gastrocnemius lateris in adults.

## Results

### Body Mass and Tibial length

As adults, spastic mice had 25% lower body mass compared to WT controls (p<0.05). Female WT mice had 31% lower body mass, compared to male WT mice, and female spastic mice had 17% lower mass than male spastic mice (p<0.05). Tibia length of spastic mice was 4.5% shorter than that of WT mice (p<0.01), and tibia of female spastic mice was 3% shorter than that of male spastic mice (p<0.05; Table 1)

**Table 1.**
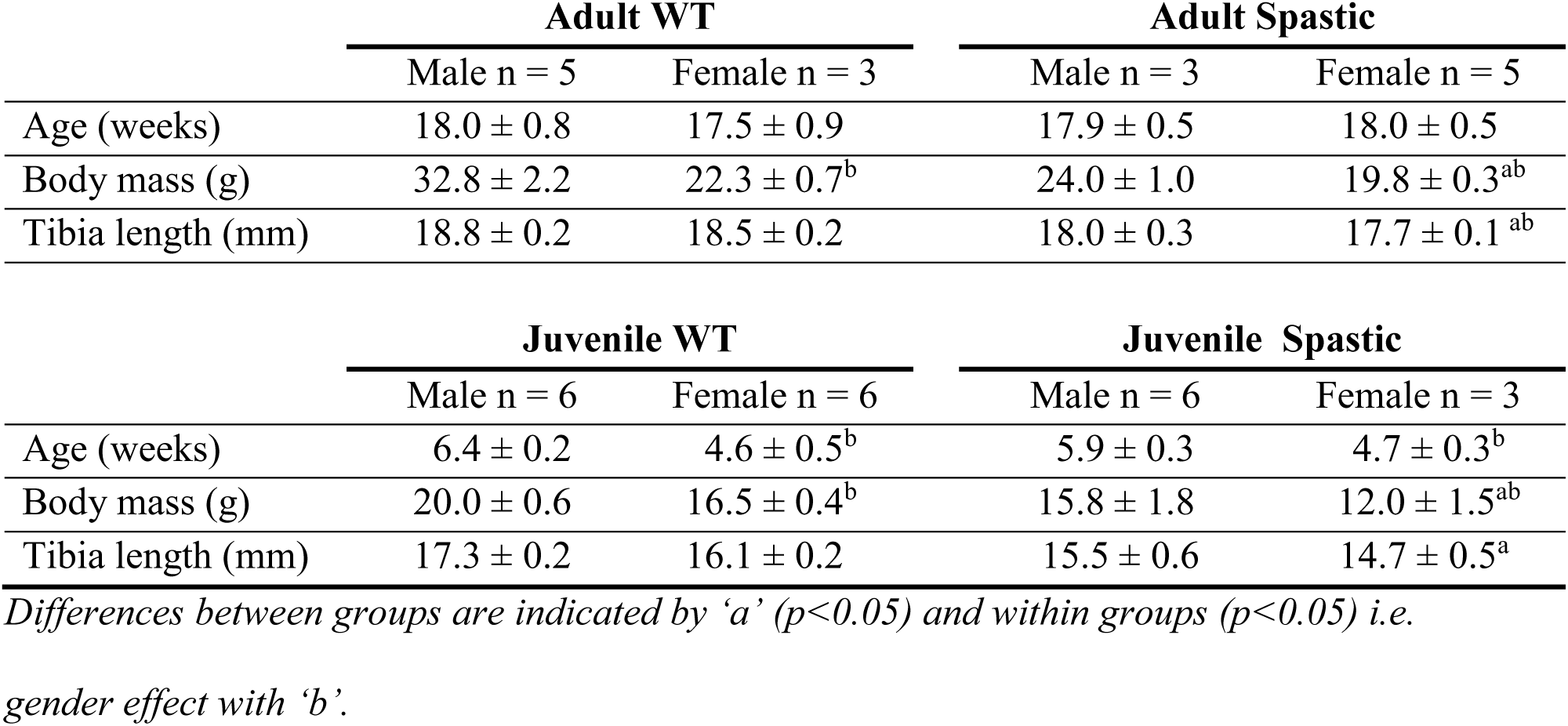
Body measurements of the adult and juvenile mice.

As juvenile, spastic mice had 20% lower body mass compared to WT controls (p<0.05). Moreover, of the juvenile mice, female mice had about 16% lower body mass, both in the female in WT and spastic group (p<0.05). Tibia length of juvenile spastic mice was 9% shorter than that of juvenile WT mice (p<0.05; Table 1).

### Motor function

Table 2 shows an overview of parameters associated with motor function performance. Activity of spastic mice on rotarod was characterized by 32% fewer rotations per minute in comparison to WT controls (p<0.001; Fig. 1A). For both genotypes, performance (i.e. rotations per minute) decreased as mice became older (p<0.0001). Grip force of both genotypes increased as mice became older (p<0.0001), however, grip force of spastic mice was 23% lower than that of WT mice (p<0.001; Fig. 1B).

**Table 2.**
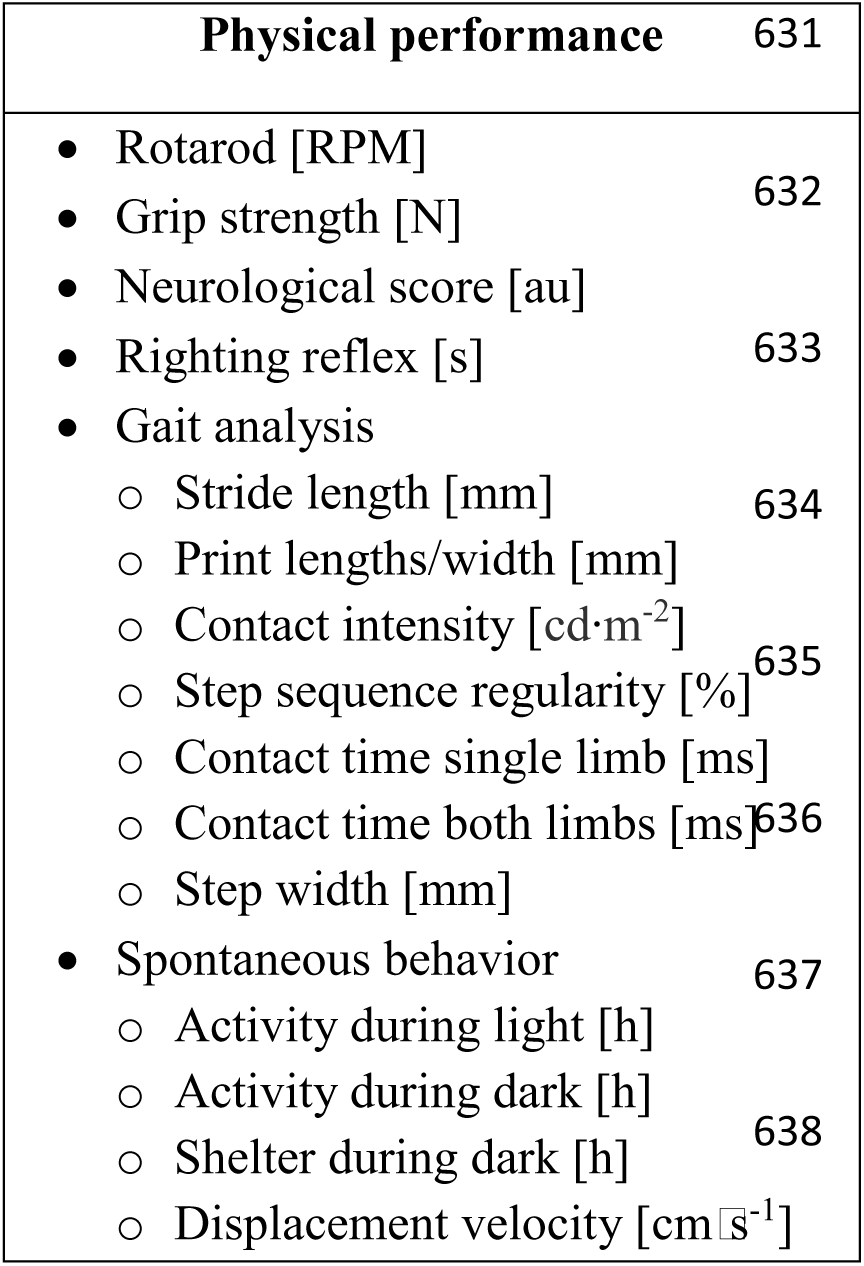
Variables reported in the physical performance result section

**Figure 1.**
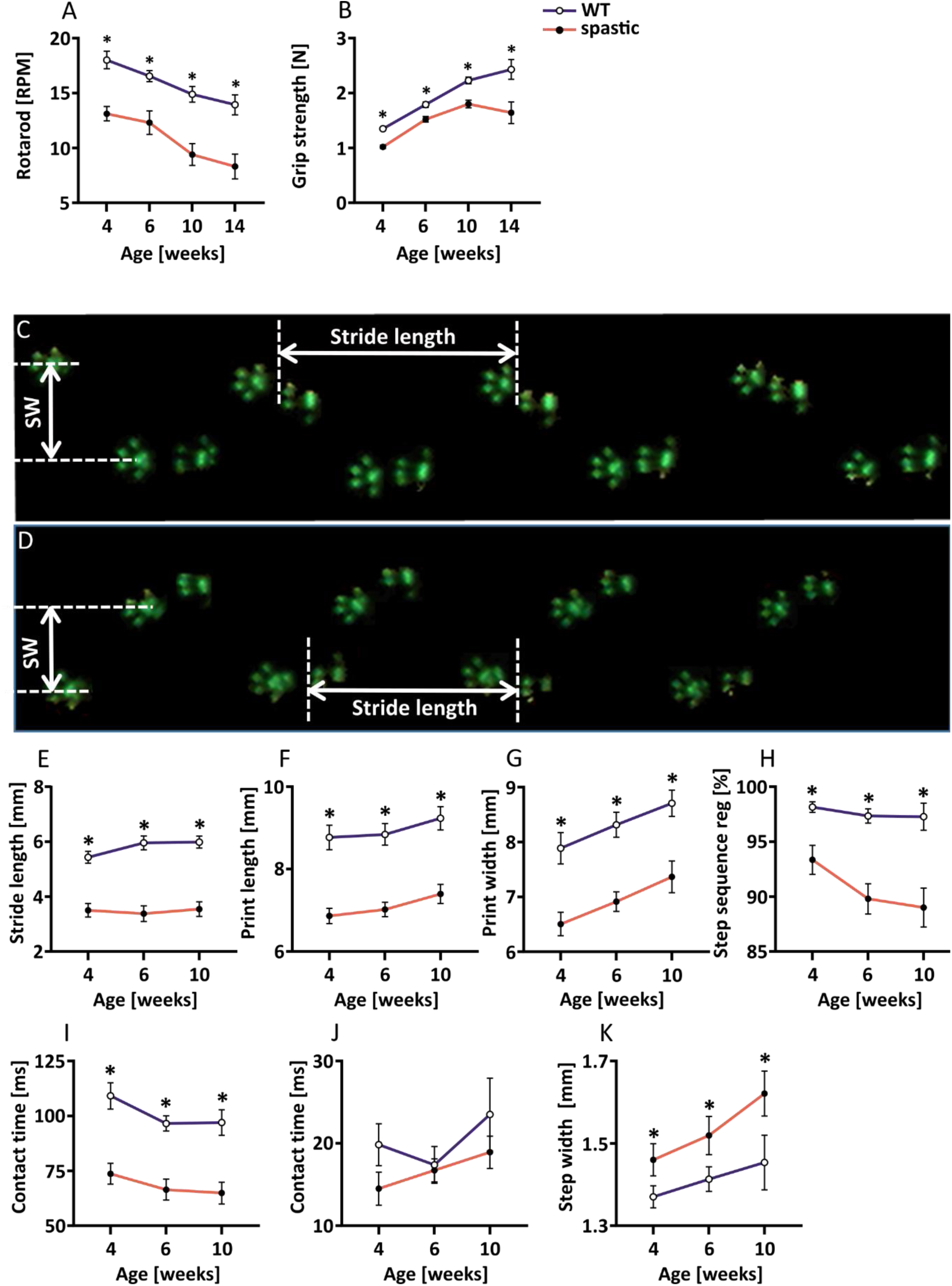
Motor function of spastic mice. (A) Rotarod performance, (B) Grip strength of front and hind limbs together. (C) Footprints example as registered by the Catwalk in WT and (D) spastic mice, (E) Stride length, (F) Print length of hind limb, (G) Print width of hind limbs, (H) Step sequence regularity index, (I) Contact time of single limb, (J) Contact time of both hind limbs together (K) Step width of front limbs.

At all ages, the neurological score of spastic mice was different from zero (i.e. score of WT mice), even though it was constant during development (Table 3). Righting reflex of spastic mice was attenuated while during growth WT mice maintained their righting time below 1 second (p<0.05; Table 3). During growth, the time to righting of spastic male mice increased by 0.78 s per week, while in spastic female time to righting increased by 0.94 s per week.

**Table 3.**
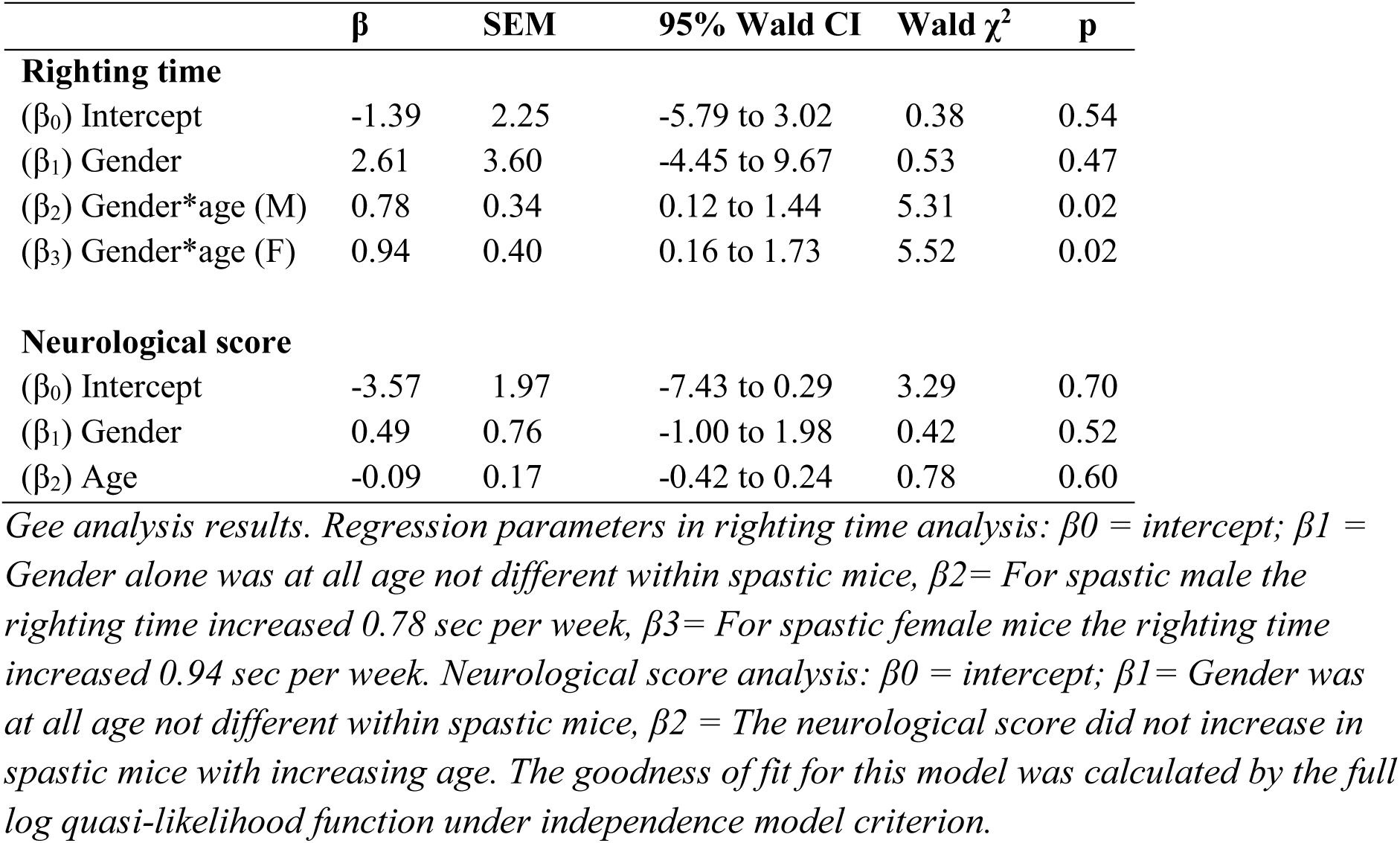
Results of righting time and Neurological Score of spastic mice

### Gait

At all ages, gait of spastic mice was characterized by shorter stride lengths, smaller print-length and width (p<0.001) together with a lower contact intensity (p<0.01; Figs. 1E to 1G). Spastic mice showed an irregular order of front and hind-limb placements (p<0.001; Fig. 1H). Compared to WT mice, spastic mice showed shorter ground contact times of single limbs (p<0.001), but unaffected ground contact times of both limbs together (Figs. 1I and 1J). With increasing age, step with of the front paws increased in both WT and spastic mice, however step with of spastic mice was substantially larger than of WT mice. (p<0.001; Fig. 1K).

### Spontaneous physical activity in home cage

During the light phase, spastic mice showed a trend towards reduced physical movement (Table 4). In contrast, in all age-groups, physical activity of spastic mice during the dark phase was 48% lower in comparison to WT controls (p<0.05; Table 4). Activity time of WT mice progressively increased during growth (p<0.0001), while after 10 weeks of age spastic mice did not increase their activity time during the dark phase. This latter was accomplished by the fact spastic mice spent 36% more time in shelter during the dark phase that compared to WT mice (p<0.01; Table 4). During development, shelter behavior of WT mice decreased 42% (p<0.0001), while that of spastic mice did not. Moreover, displacement velocity during activity bouts of spastic mice was between 22% lower than that of WT mice (p<0.0001; Table 4).

**Table 4.**
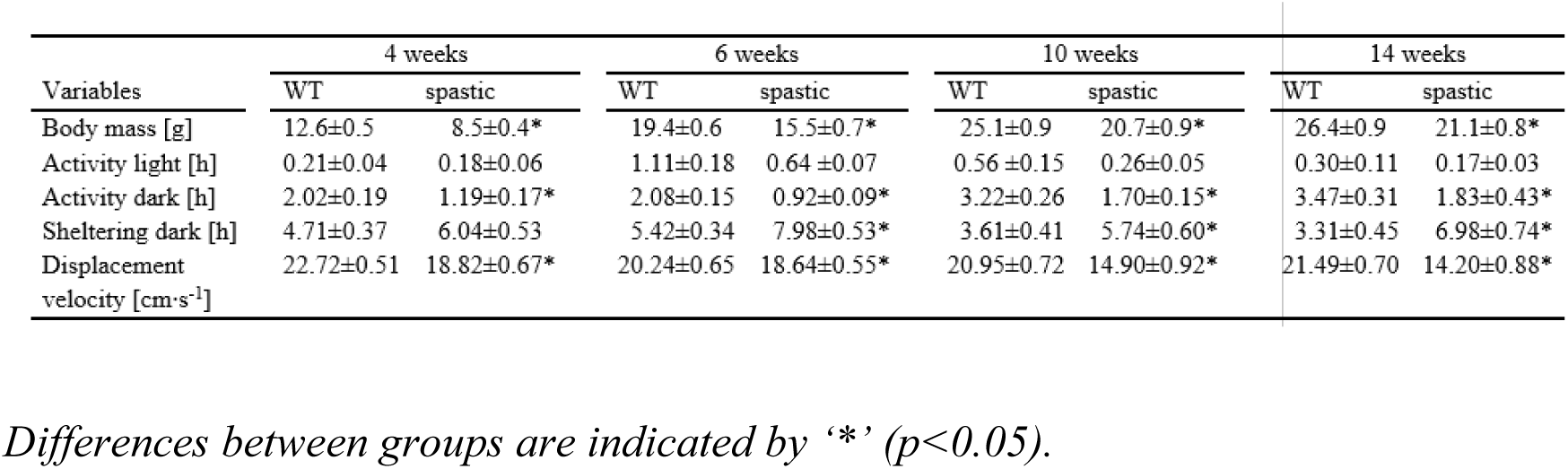
Variables of Body mass, and behavior during growth measured in automated home-cage.

#### Contractile force characteristics of SO and GM muscles

Table 5 shows contractile force characteristics, morphological and histological determinants as well as parameters of muscle growth and degradation.

**Table 5.**
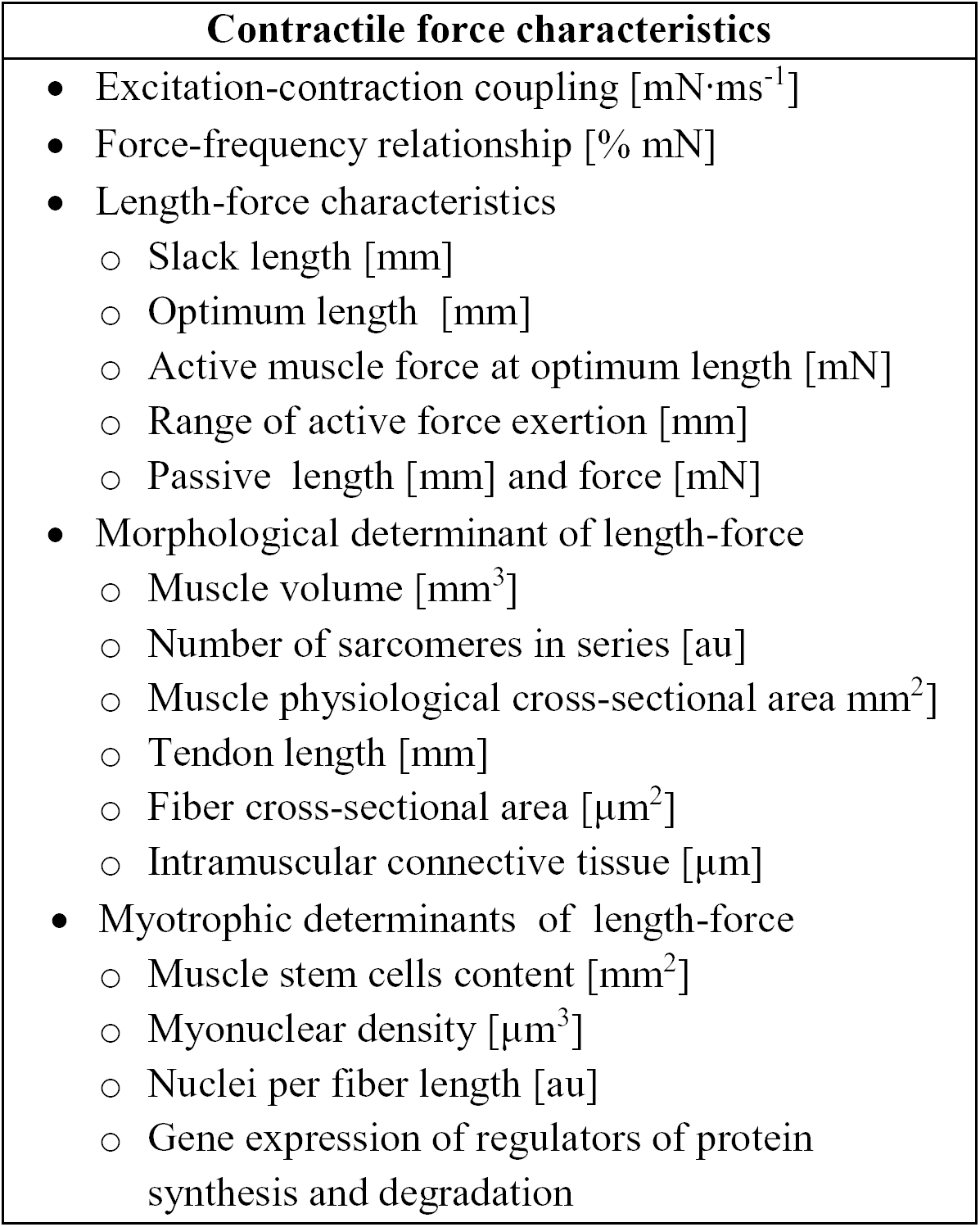
Variables reported in the contractile force characteristics and length-force determinants sections

### Excitation-contraction (E-C) coupling

In adult mice, the maximal rate of force development of SO and GM did not differ between spastic and WT animals. For GM, absolute maximal rates of force development of female vs male mice spastic mice were 48 and 30% lower, respectively (p<0.05; Figs. 2A to 2D).

**Figure 2.**
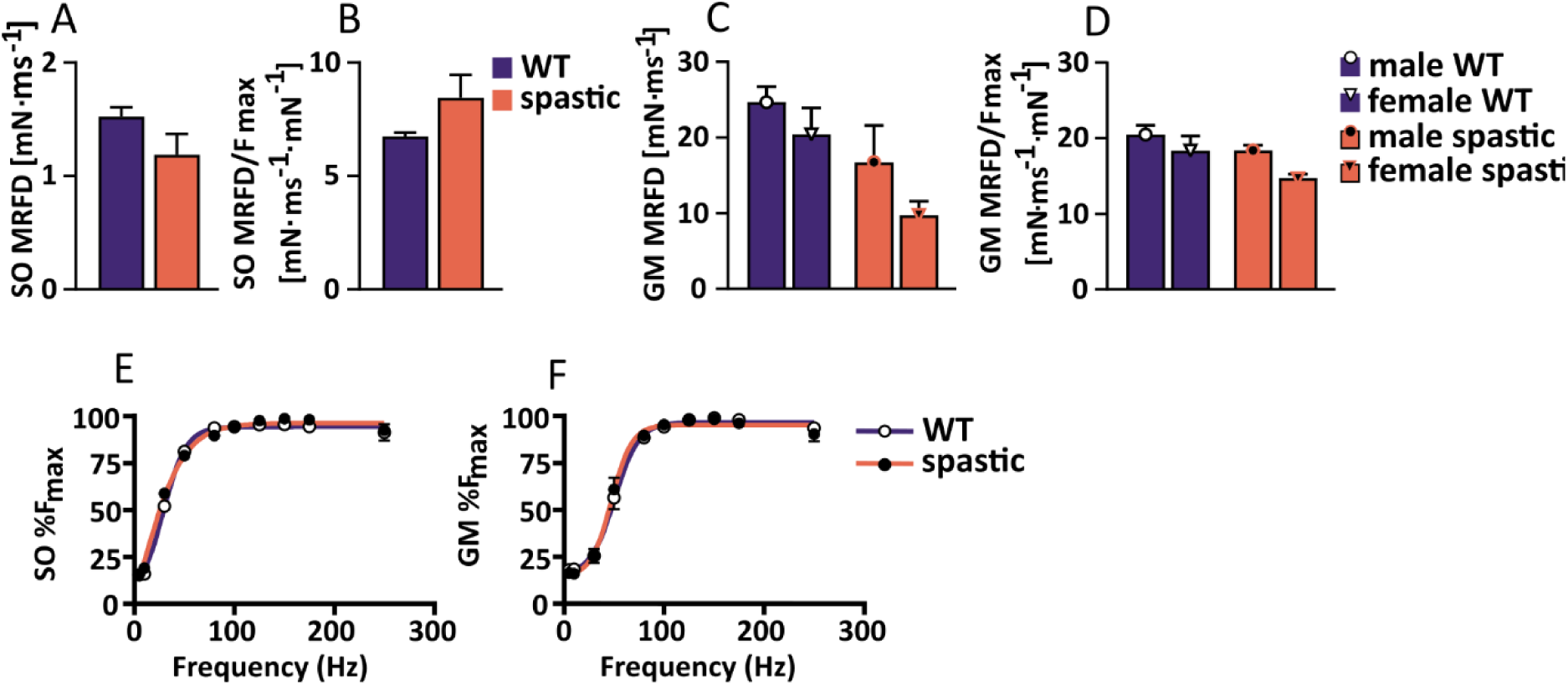
Effects of spasticity on maximal rate of force development and force-frequency relationship in adult. (A) Absolute rate of maximal force development of SO muscle, (B) Rate of maximal force development of SO normalized for maximal force, (C) Absolute rate of maximal force development of GM muscle, (D) Rate of maximal force development of GM, (E) Force-frequency relationship of SO muscle, (F) Force-frequency relationship of GM muscle.

For juvenile SO, no difference could be shown between spastic and WT animals. For GM muscle, male WT mice showed a 47% higher maximal rate of force development and 30% higher normalized maximal rate of force development than spastic male mice (p<0.05; Supplementary Figs. 1A to 1D).

### Force-frequency relationship

Active forces of SO and GM as function of stimulation frequency did not differ between spastic and wild type mice neither in adults nor in juveniles (Figs. 2E to 2F and Supplementary Figs 1E to 1F).

### Length-force characteristics of SO and GM muscles

In adult mice, at the ascending limb of the SO length-force curve, spastic active SO force was 13% lower compared to that of WT SO (p<0.05). Force of the active muscle at optimum length was 34% lower, indicating that spastic SO was substantially weaker than WT SO (Fig. 3A). Muscle lengths at which force was actively exerted were similar in both groups (Fig. 3B).

**Figure 3.**
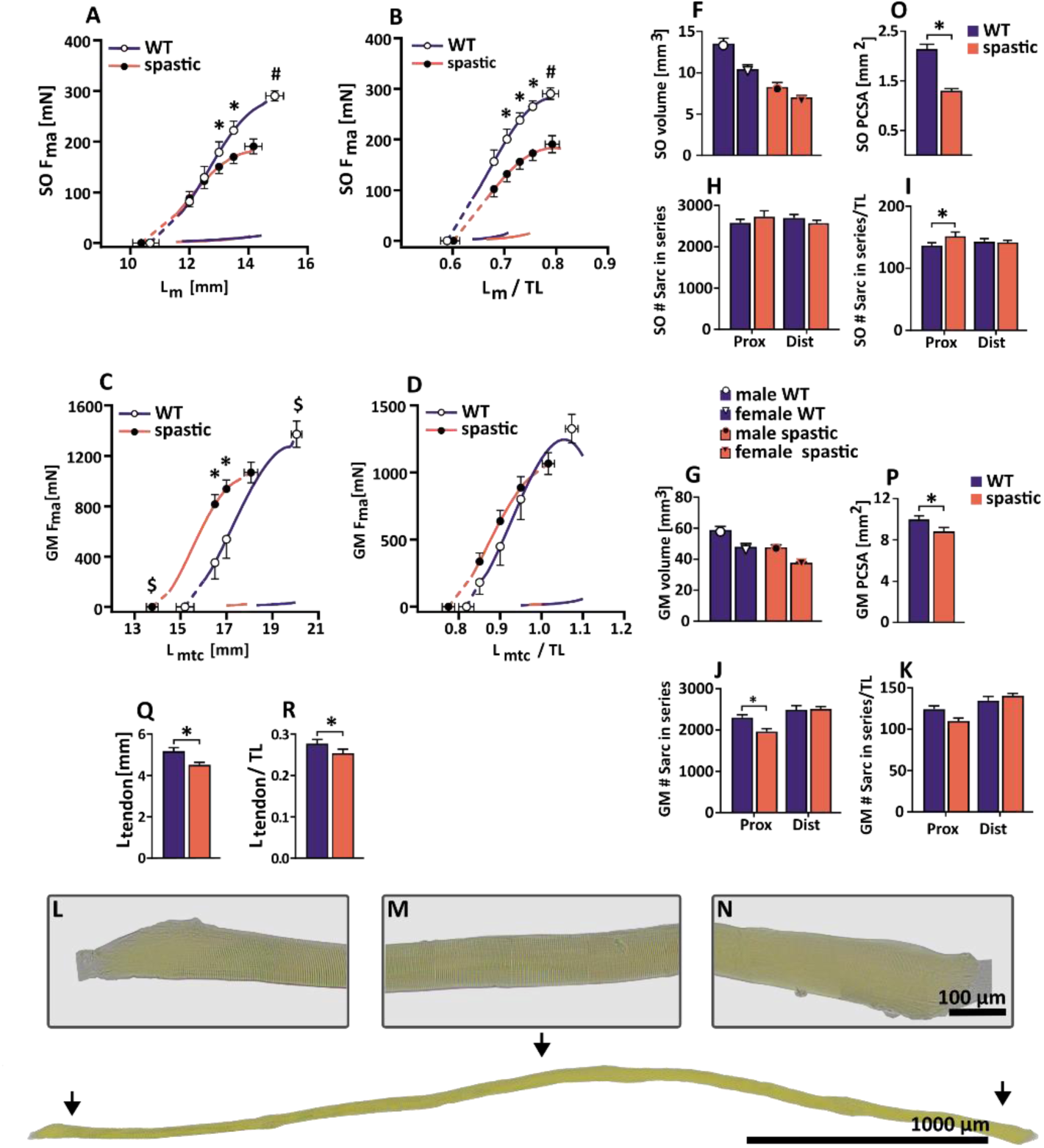
Effects of spasticity on adult muscle length-force relationship as well as determinants of muscle length and muscle force. (A) SO absolute length-force relationship, (B) SO length-force normalized for tibia length, (C) GM absolute length-force relationship, (D) GM length-force relationship normalized for tibia length, (E) Typical example twitch and tetanus at optimum muscle length of WT (blue) and spastic (red) SO muscle, (F) SO muscle volume, (G) GM muscle volume, (H) SO number of sarcomeres in series of proximal (Prox) and distal (Dist) fibers, (I) SO number of sarcomeres in series of Prox and Dist fibers per unit of tibia length, (J) GM number of sarcomeres in series of Prox and distal Dist fibers, (K) GM number of sarcomeres in series of Prox and Dist fibers per unit of tibia length, (L-N) An example of a muscle fiber with visible serial sarcomeres, the arrangement of actin and myosin filaments are clearly visible as light (I-band) and dark (A-band) cross striation. At the beginning and end of the fiber, the finger-like invaginations are shown. (O) SO physiological cross-sectional area (PCSA), (P) GM PCSA, (Q) Achilles tendon length, (R) Achilles tendon length normalized for tibia length.

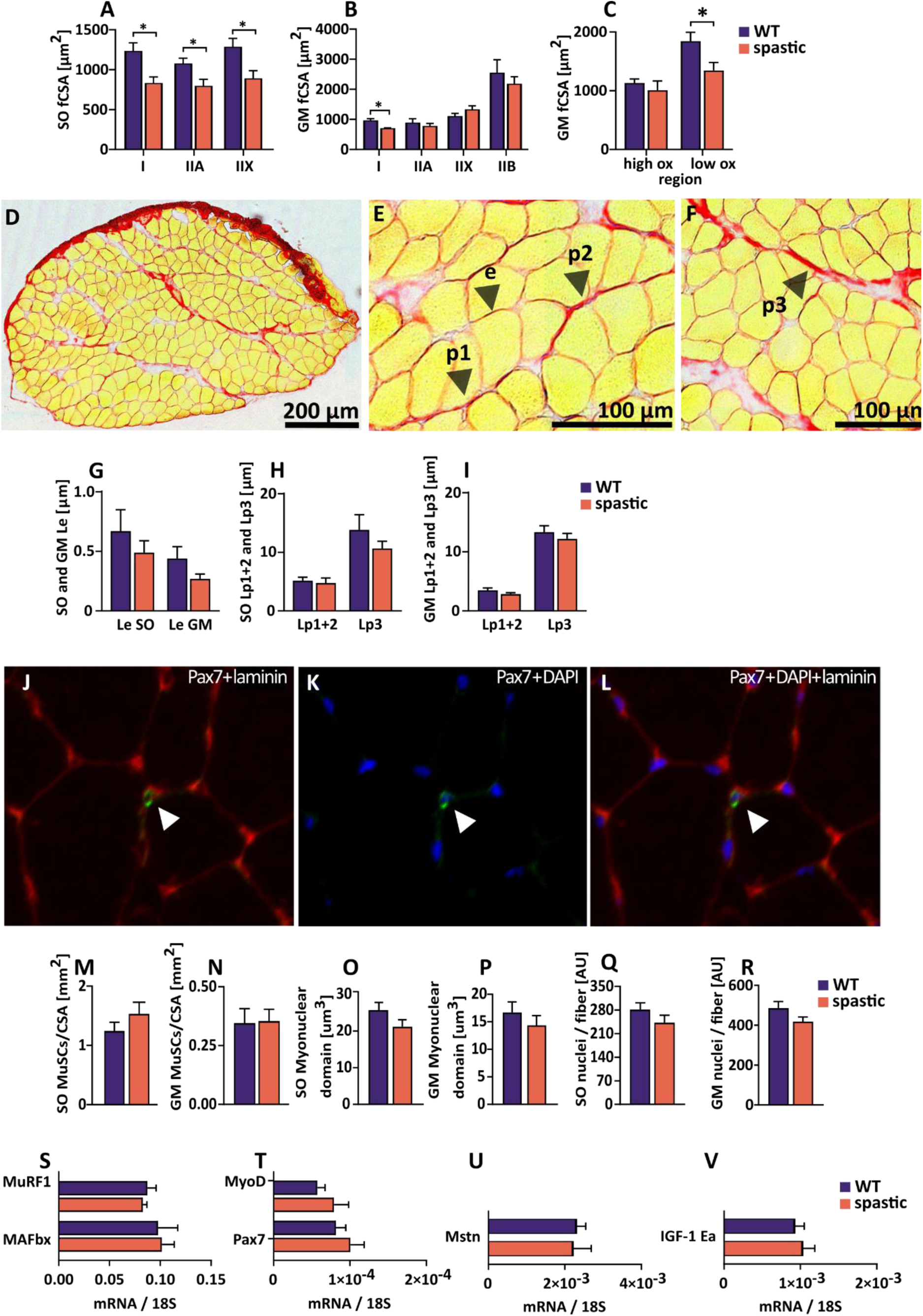

At the ascending limb of the GM length-force curve, spastic GM force was 49% higher compared to that of WT animals, while force of the active muscle at optimum length was 22% lower (p<0.05). In spastic GM, active slack and optimum lengths were about 10% shorter than those of WT GM, indicating that for GM the length-force curve was shifted to shorter lengths (p<0.05; Fig. 3C). This implicates that at similar lengths, spastic muscles exerted higher forces than WT muscles. By normalizing muscle-tendon complex length to tibia length, spastic GM length-force curve shifted towards the curve of WT GM, and absolute forces of both groups were similar (Fig. 3D). Muscle-tendon complex length range over which spastic GM was able to exert active force did not differ from that of WT muscle. Moreover, active slack and optimum length were not statistically different after normalization, neither was the range of active force exertion (Table 6).

**Table 6.**
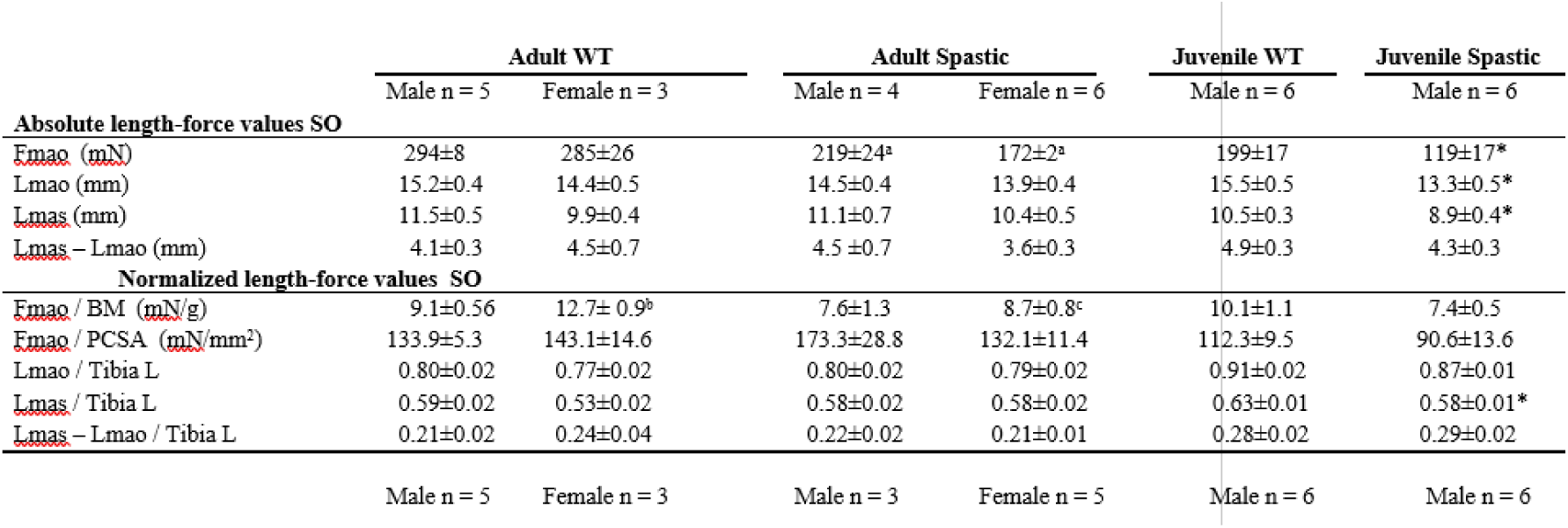
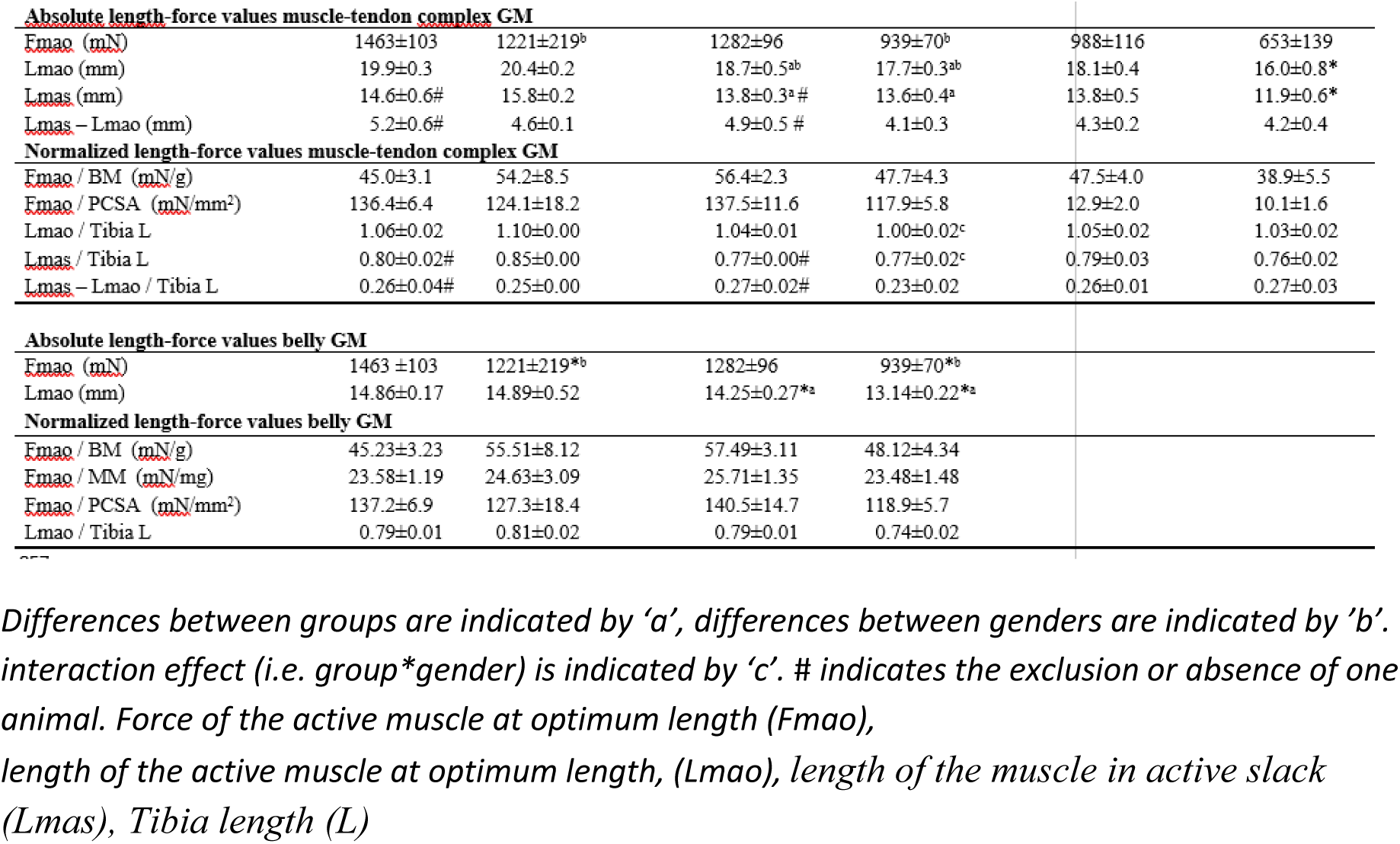
Adult and juvenile length-force relationships variables of SO and GM muscles

In juvenile spastic mice, absolute SO forces at the ascending limb of length-force curve were 57% higher (p<0.05) than those of WT juvenile mice, while force at optimum muscle length was 40% lower (p<0.05). In spastic SO, active slack and optimum muscle lengths were, respectively, 15% and 14% lower compared to those of WT SO (p<0.05; supplementary results). Normalization of muscle length by tibia length yielded similar curves for both groups, although for spastic SO active slack length remained 8% shorter than for WT SO (p<0.05; Supplementary Figs. 2A and 2B and Table 6).

For juvenile spastic GM, length-force curves showed substantial variation in length range of active force exertion. Therefore, it was not possible to plot them together in one figure and calculate mean curves. Absolute values at slack and optimum lengths showed that optimal muscle force of spastic juvenile GM was similar to that of WT GM, while active slack length optimum length were 14% and 12% shorter, respectively (p<0.05). Normalization of active slack length, and optimum length range of force exertion by tibia length did not reveal any differences between groups (Supplementary Figs. 2C).

### Morphological determinants of muscle length-force characteristics

#### Muscle volume

In spastic adult mice, SO volume was 36% smaller compared to that of WT mice (p<0.05). Adult male mice had 20% greater SO volume compared to that of female mice (p<0.001; Fig. 3F). Similar results were shown for GM volume (i.e. in spastic mice 18% smaller than in WT mice). Note, that in both groups, for females GM volume was 18% smaller than for male mice (p<0.001; Fig 3G).

SO muscle volume of spastic juvenile male mice was 36% smaller (p<0.01) and GM volume was 28% smaller (p<0.05) than that of WT juvenile mice (Supplementary Figs. 2D and 2E).

#### Number of sarcomeres in series

For adult mice, the number of sarcomeres in series of SO in both proximal and distal regions were similar in spastic and WT mice. After normalization for tibia length, the number of sarcomeres in the proximal region of spastic SO was higher than that in the same region of WT SO (p<0.01; Fig. 3H). In addition, normalization yielded a difference between genders, WT and spastic female mice had fewer sarcomeres in series within SO compared to male WT and spastic mice (p<0.05; Fig. 3I). For spastic male GM, the number of sarcomeres in series in the proximal region was slightly lower than in the WT (p<0.05; Fig. 3J). After normalization for tibia length these differences disappeared (Fig. 3K).

For juvenile mice, the number of sarcomeres in series of SO and GM in both proximal and distal regions were similar in spastic and WT mice (Supplementary Figs 2F to 2I).

#### Muscle physiological cross-sectional area

For adult mice, physiological cross-sectional area values of spastic SO and GM were 39 and 14% smaller compared to those of WT mice, respectively (p<0.05; Figs. 3O and 3P).

In juvenile spastic male mice, SO physiological cross-sectional area was 26% smaller than in juvenile WT mice (p<0.05), however, physiological cross-sectional area of GM did not differ from that in WT mice (Supplementary Figs. 2J and 2K,).

#### Tendon length

For adult spastic mice, Achilles tendon length at GM optimum length was 13% shorter (p<0.01) compared to that of WT mice (Fig. 3Q) and 10% shorter when normalized for tibia length (p<0.05, Fig. 3R).

In juvenile spastic male mice, Achilles tendon length was 24% shorter compared to that of WT mice (p<0.05), but normalized Achilles tendon length was similar for spastic and WT mice (Supplementary Figs. 2L and 2M).

#### Fiber cross-sectional area

In adult spastic SO, for all muscle fiber types, the fiber cross-sectional area was smaller compared to that of their WT litter mates (i.e. 32% in type I, 25% in type IIA and 31% in type IIX, respectively (p<0.05; Fig. 4A).

In spastic adult GM, cross-sectional area of type I muscle fibers was 27% smaller than that in WT GM (p<0.05; Fig 4B). However, the cross-sectional area of type IIA, IIX and IIB were similar between both groups. In addition, the total cross-sectional area of fibers in the lox oxidative muscle region (consisting mainly of type IIB and IIX fibers) in spastic adult GM was 41% smaller than that in WT GM (p<0.05; Fig. 4C). Moreover, the total cross-sectional area of fibers in the high oxidative muscle region was similar between both groups.

In juvenile spastic SO, fiber cross-sectional area of type I fibers were 19% smaller compared to that in juvenile WT muscle (p<0.05; Supplementary Fig. 3A), although in both SO and GM a trend was shown towards a smaller fiber cross-sectional area in juvenile spastic fibers, no significant differences were shown for fiber cross-sectional area of other muscle fiber types (supplementary Figs. 3A to 3C).

#### Intramuscular connective tissue content

Figures 4G to 4I show for both spastic and WT the content of endomysium and perimysium in adult muscles. Results for juvenile SO and GM muscles are presented in supplementary Figs. 3D to 3F. For both ages, mean endomysium and perimysium thickness did not differ, there were also no differences between spastic and WT mice in the fraction of endomysium per unit cross-sectional area, as well as in the amount of endomysium per muscle fiber.

#### Myotrophic determinants

The above results show that spastic plantar flexor muscles are smaller in volumes and physiological cross-section area. To obtain insight in the mechanisms underlying the attenuate muscle growth, muscle stem cell and myonuclear densities were analysed as well as expression of myogenic regulatory factors, growth factors and enzymes involved in degradation of muscle protein.

In both adult and juvenile mice, the number of Pax7 stained muscle stem cells per cross-section area in SO and GM did not differ between WT and spastic muscles (Figs. 4M and 4N and supplementary 3G and 3J).

In adult spastic SO and GM, numbers of myonuclei per myofiber did not differ from those in WT mice (Fig. 4). The myonuclear domain in SO and GM also did not differ between both groups (Figs. 4O and 4P). Results from juvenile muscles show that the lack of difference between spastic and WT muscle is not age dependent (Supplementary Figs. 3G to 3L). The smaller physiological cross-sectional area in spastic mice could therefore not be explained by a reduced density of muscle stem cells myonuclei.

Figures 4Q to 4T show for gastrocnemius lateralis mRNA levels of, growth factors (IGF-1Ea and myostatin) and E3-ligases, (i.e. Murf1 and MAFbx), involved in degradation of muscle protein. For none of these genes, expression levels differed between spastic and WT muscle. These observations indicate that morphological differences between WT and spastic muscles are likely not explained by elevated expression of genes involved in muscle protein degradation nor by a diminished growth factor expression.

#### Determinants for muscle endurance

#### Muscle fatigue resistance and determinants of endurance capacity

Table 7 shows an overview of parameters determined in the endurance capacity and fatigue resistance.

**Table 7.**
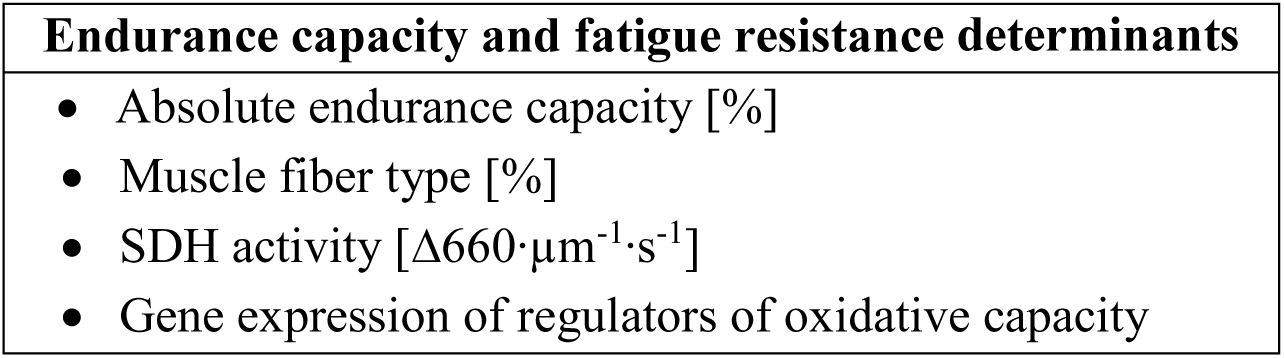
Variables reported in the endurance capacity and fatigue resistance determinants section

In adult mice, fatigue resistance was not affected in spastic SO, as during the 2 minutes of stimulation the percentage decrease in force of spastic SO was not different compared to that of WT SO (Figs. 5A and 5B).

**Figure 5.**
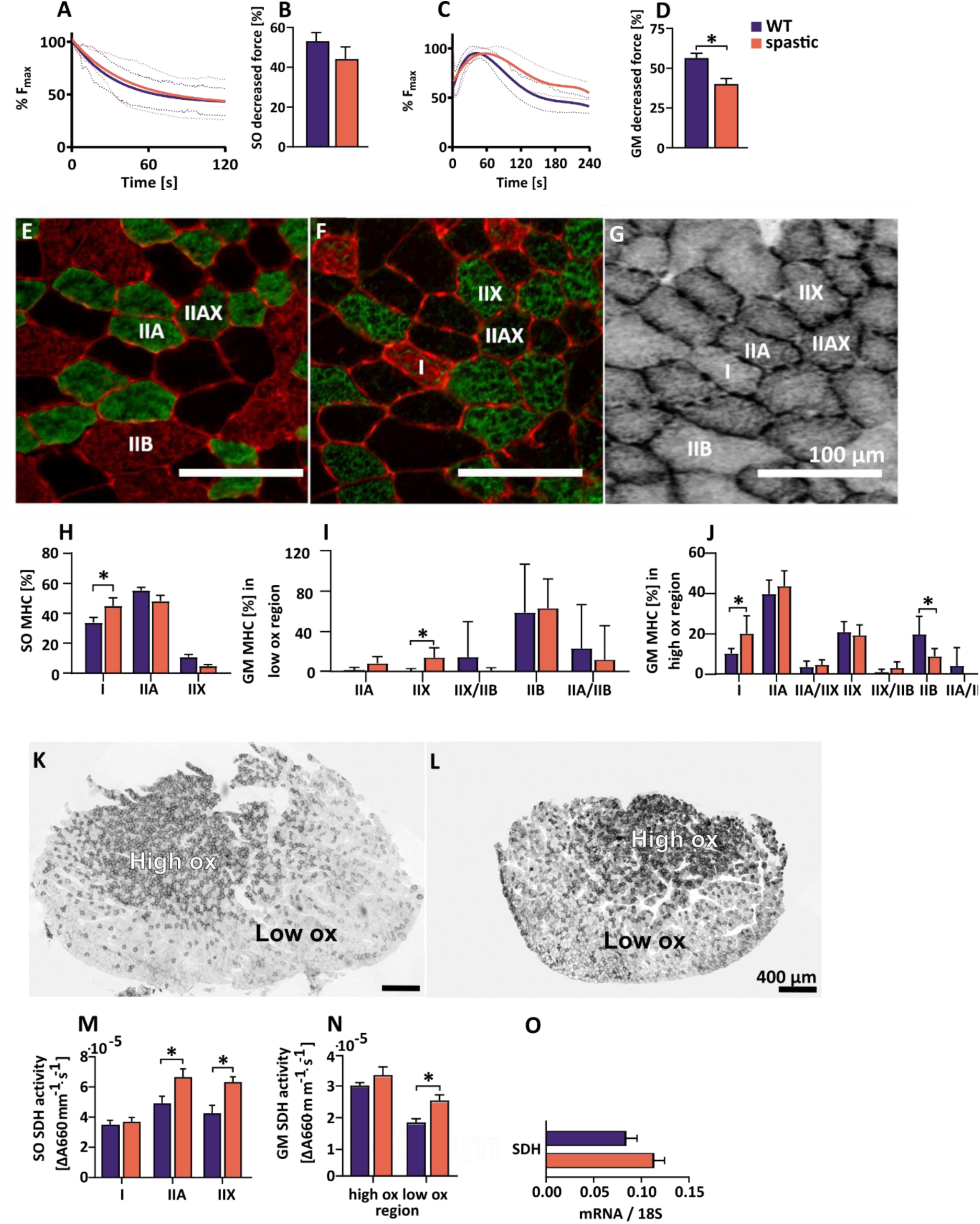
Endurance capacity of adult SO and GM as well as endurance determinants. (A) SO endurance curve, (B) SO percentage of decreased force at the end of the fatigue protocol, GM endurance curve, (D) GM percentage of decreased force at the end of the fatigue protocol, (E-G) Typical example of muscle fiber type matching with SDH stained cross-section. (H) SO MHC distribution, (I) GM MHC distribution in the low oxidative muscle region, (J) GM MHC distribution in the high oxidative muscle region, (K-L) Typical examples of WT and spastic GM muscles, respectively. (M) SO SDH activity, (N) GM SDH activity, (O) gene expression factors involved in the regulation of muscle metabolism, expression factors are relative to the expression of 18S RNA.

For GM, the decrease in maximal force of spastic GM after 4 minutes of stimulation was 16% lower than those of WT mice (p<0.01; Figs. 5C and 5D). These results show that spastic GM was more fatigue resistant than WT GM.

In juvenile mice the decreases in maximal force of SO and GM were similar between groups. Even though endurance curves for SO and GM showed later fatigue onset in spastic muscles, total decrease in force was not different between groups (Supplementary Figs. 4A to 4D).

#### Muscle fiber typing

Myosin heavy chain staining was used to compare muscle fiber type distributions. In adult spastic SO, the type I percentages was 27% higher than in WT SO (p<0.01; Fig. 5H). The higher percentage of type I MHC in spastic SO was mainly caused by a gender effect. Spastic adult female mice had 30% higher percentages of type I MHC compared to spastic adult male mice, but also 34% higher than WT female (p<0.001), which caused a difference of 27% between spastic and WT SO.

For GM muscle, the percentage of type IIX within the low oxidative region, of spastic mice was higher than that of WT GM (p<0.01; Fig. 5I). Within the high oxidative region of spastic muscle, the percentage of type I fibers was 46% higher than within this region of WT muscles (p<0.05). Within the high oxidative region, WT muscles showed 55% higher percentages of type IIB fibers (p<0.05; Fig. 5J) than spastic muscles.

In juvenile SO, muscle fiber typing showed similar distributions in both groups (Supplementary Fig. 4E). In the low oxidative region of juvenile GM, the presence of hybrid muscle fibers was below 1%, whereas in the high oxidative region IIA/IIX hybrid muscle fibers were present. In the high oxidative muscle region of spastic juvenile GM, 14% less type IIB muscle fibers were shown than in the same region in juvenile WT GM (p<0.05; Supplementary Figs 4F and 4G).

#### Succinate dehydrogenase activity (SDH)

In adult spastic SO, SDH-activity in type IIA and IIX muscle fibers was higher than in that these fibers in WT SO (p<0.05; Fig. 5M), which indicates that in particular within spastic, fast type SO muscle fibers the oxidative capacity was higher than in WT SO fibers.

For adult mice, SDH-activity of muscle fibers within the low oxidative region of spastic GM were 27% higher than that of same type of fibers within WT GM (p<0.01; Fig. 5N). SDH activity in the high oxidative muscle region of GM was similar in WT and spastic mice.

SDH activity in type I, IIA and IIX fibers in spastic juvenile SO was higher than that in similar fiber types in WT SO (p<0.05; Supplementary Fig. 4H). GM SDH activity did not differ between juvenile WT and spastic mice (Supplementary Fig 4I). Figure 5O shows for the gastrocnemius lateralis mRNA levels for SDH, expressed relative to 18S RNA levels. SDH mRNA levels were similar in spastic and WT mice.

## Discussion

The aim of this study was to quantify in detail motor function, gait and physical activity of spastic and WT mice as well as to characterize their calf muscles, SO and GM, with respect to their contractile force characteristic, morphology and histological phenotype at both juvenile to adult age. These two plantar flexor muscles were chosen, as their geometry and function differ from each other. In rodents, the SO is a parallel fiberd muscle and its main function is to control posture, whereas the GM is a highly pennate muscle mainly involved in the push off during locomotion. Results of the present study are discussed and compared to those of human studies, to obtain insight in whether spasticity affects growth, as well as physical performance and muscle function and phenotype of humans and mice with spasticity in similar way.

### Effects of spasticity on growth

Here we show that growth of spastic mice is attenuated (i.e. lower body mass at juvenile age and shorter tibia length at juvenile and adult age). Children with spasticity also show attenuated growth which is manifested in a lower body weight and a shorter stature (Shapiro et al. 1986; Kuperminc et al. 2009; Stevenson et al. 2006; Day et al. 2007). Several factors have been suggested to be responsible for the growth deficit (i.e. malnutrition, growth hormones deficiency, low level of physical activity and intrinsic genetic factors). Impaired growth in children with spasticity has been associated with malnutrition due to feeding difficulties and diminished serum levels of growth hormones (Shapiro et al. 1986; Stallings et al. 1993; Coniglio et al. 1996). However, low levels of physical activity may also contribute to a reduced stimulus for growth. In mice, the observed attenuated growth may also be due to a low level of physical activity. However, the contribution of malnutrition and intrinsic factors cannot be excluded. As muscle mRNA expression levels of factors associated with protein synthesis and degradation in spastic muscles were similar to those of controls, the attenuated growth of spastic mice and their muscles is likely not caused by an altered regulation of protein synthesis, nor by increased degradation. Children with spasticity and spastic mice may experience reduced stimulus for growth due to low levels of physical activity.

### Effects of spasticity on physical performance

Spastic mice showed impaired motor function, altered gait, and lower physical activity. During walking, spastic mice show a typical hopping pattern, characterized by shorter stride lengths, less and irregular surface-paw contact as well as increased base of support area of front paws. These gait characteristics indicate that in spastic mice, the coordination and balance control are hampered. Spastic mice showed also high neurological scores and an impaired righting reflex when placed on their back. A lower gait velocity and shorter step length are also characteristic for gait of children with spasticity (Van Den Hecke et al. 2007). In addition, these spastic children show lower dynamic balance capacity compared to typical developing children (Jeng, Liao & Hou, 1997). In children with spasticity, muscles of the lower extremities are weaker than in typical developing children. (Wiley and Damiano 1998). Moreover, for children with spasticity show a reduced travel distance during a 5 minutes walking test (Eken et al. 2019).Therefore, the energy demand of lower leg muscles during walking is high, which results in an early onset of fatigue (Parent et al. 2016). In children with spasticity, impaired mobility due to a hampered coordination in combination with a reduced exercise tolerance implicates participation restrictions and activity limitations. As a consequence total daily physical activity of these children is reduced compared to that of typical developing children (Beckung and Hagberg 2002; Bjornson et al. 2007b; Zwier et al. 2010; Oftedal et al. 2015; Keawutan et al. 2017). Although for mice the exact reason for their observed low physical activity is not clear, the impact on muscle development of both spastic mice and spastic children seems similar.

### Effects of spasticity on muscle length range of force exertion

The length range over which a muscle is able to generate force is determined by several factors: The length of myofilaments, the number of sarcomeres in series, number and length distribution of sarcomeres within muscle fibers, change in fiber and aponeurosis angle with change in muscle length, as well as the length and mechanical properties of tendinous structures within the muscle-tendon complex. Another determinant to account for is the fiber cross-sectional area. In pennate muscle, muscle belly length is also determined the length component of the physiological cross-sectional area (Huijing 1998; Bénard et al. 2011; Lieber and Bodine-fowler 1993; Weide et al. 2015).

For SO, a parallel-fibered muscle, absolute and normalized length range of active force exertion did not differ between spastic and WT mice. This lack of differences is in line with the observed lack of differences in the number of sarcomeres in series between spastic and WT SO. In spastic rats, length-force characteristics of SO muscles were also not different from control muscles (Olesen et al. 2014). These results show that in rodent SO muscle, the length range of active force exertion is not affected by spasticity. In children with spasticity it is unknown how SO the muscle length differs from that of typical developing children.

For the spastic GM muscle-tendon complex, the absolute length range of active force exertion was shifted towards shorter lengths, while length range of active force exertion was similar. This shift by a shorter Achilles tendon, a lower number of sarcomeres in series of the proximal muscle region and a smaller physiological cross-sectional area. After normalization by tibia length, muscle-tendon complex lengths were similar between spastic and WT GM. This effect of normalization is in line with similar normalized serial sarcomere numbers between spastic and WT GM, which indicates that the attenuated longitudinal growth of the tibia in spastic mice implicates a reduced growth stimulus for the plantar flexor muscles and Achilles tendon.

In children with spasticity, growth of spastic muscle-tendon complex is different from typical developing children. In children with spasticity, GM muscle fascicle and muscle belly lengths are lower than in typical developing children. In addition, their Achilles tendon is longer (Barber et al. 2011; Herskind et al. 2016; Malaiya et al. 2007; Mohagheghi et al. 2008; Barber et al. 2012; Gao et al. 2011; Wren et al. 2010). In children with spasticity, morphological and geometrical adaptations causing hyper-resistance of spastic muscles are highly heterogeneous, and depend on several factors such as severity and location of brain lesion, affected body part(s) (e.g. diplegic/quadriplegic), growth and intervention-history (e.g. Koman et al. 2004; Shortland et al. 2004; (Kerr Graham & Selber 2003; Davids et al. 2014). Moreover, morphological and geometrical adaptations within and between spastic muscles seem to differ in children with spasticity (c.f. Malaiya et al. 2007; chapter 5 in the thesis of G. Weide). Within the current spastic mouse model, morphological adaptations were homogenous, and muscle morphology was independent of severity of spasticity and intervention-history. Therefore, we suggest that muscle length range over which force is actively exerted is mainly affected by attenuated growth.

### Effects of spasticity on muscle force generating capacity

In this study we show that the force generating capacity of spastic SO is lower than that of WT mice. A trend towards lower muscle forces was noticed for GM muscle, however significance could not be shown. Lower force values are explained by the smaller physiological cross-sectional area of spastic muscles. Normalization of forces by physiological cross-sectional area (yielding specific muscle force) showed that reductions in force and physiological cross-sectional area were in the same order of magnitude, which indicates that spasticity caused a radial growth deficit, rather than an impaired force generating capacity of the muscle (i.e. reduced quality). The observation that spastic muscle has a smaller physiological cross-sectional area is in line with data from children with spasticity, as spastic children in general show a reduced physiological cross-sectional area (Haberfehlner et al. 2016; Bandholm et al. 2009; Moreau et al. 2010). To the best of our knowledge, for the SO muscle of children, there is no data available regarding physiological cross-sectional area. Therefore, a direct comparison with ultrasounds studies cannot be made. For GM in children with spasticity, ultrasound studies have shown that young age calf muscle morphology is only slightly different from typical developing children, however from the age of 6 years there is a slower increase of physiological cross-sectional area (Barber et al. 2011). The present study also shows that in juvenile spastic mice, an increase in physiological cross-sectional area is less affected than in adult mice. Spastic mice and children with spasticity show similar attenuated increases in physiological cross-sectional area of spastic muscles

### Effects of spasticity on passive length force characteristics

Passive force as function of muscle length is determined by slack length and stiffness of muscle belly as well as tendon (Zuurbier and Huijing, 1991). For parallel fibered muscles, the passive slack length of the muscle belly is largely determined by the number of sarcomeres in series within the muscle fibers. For pennate fibers muscles, passive slack length is also determined by the length component of the physiological cross-sectional area. The stiffness of a muscle-tendon complex is determined by the number of sarcomeres in series, the fiber cross-sectional area (i.e. the total number of sarcomeres arranged in parallel, physiological cross-sectional area), tendon(s) length and connective tissue content and its mechanical material properties (particularly of the perimysium).

Children with spasticity show steeper relationships between ankle and knee joint angles and external applied moments compared to typical developing children (Haberfehlner et al. 2016; Bénard et al. 2010). This means that in children with spasticity, the resistance to joint extension is higher. Muscle-tendon complex stiffness in children with spasticity is likely not caused by an increased number of parallel arranged sarcomeres as the physiological cross-sectional area of spastic muscles is generally smaller, however it is possible that a reduced number of sarcomeres in series, contributes to muscle stiffness. Another factor that could have contributed to the enhanced muscle stiffness, is the tendon length. However, several studies have shown increased tendon lengths in children with spasticity, while similar tendon lengths in typical developing children and children with spasticity have been reported as well (c.f. Barber et al. 2011; Herskind et al. 2016; Malaiya et al. 2007; Mohagheghi et al. 2008; Barber et al. 2012; Gao et al. 2011; Wren et al. 2010; Haberfehlner et al. 2016). This indicates that in spastic muscle-tendon complex, the tendon likely does not contribute to the increased muscle stiffness. Moreover, connective tissue content has been shown to be increased in some muscles of some children with spasticity, however large variations between muscles and subjects does exist (e.g. De Bruin et al. 2014; Smith et al. 2011, booth et al 2001). A possible explanation for this could be the severity of spasticity as this seems to be related to the amount of connective tissue content y (Booth, et al. 2001).

In SO of spastic mice, there were no differences in passive length-force characteristics. The number of sarcomeres in series in normalized SO did not differ between spastic and WT mice, nor did the connective tissue content. The smaller physiological cross-sectional area and lower optimal force in spastic SO indicate that the total number of sarcomeres arranged in parallel was lower. Based on these results, one would expect a reduced stiffness in spastic SO, however this could not be shown. The lack of difference in muscle stiffness between spastic and WT SO could be explained by spastic muscles having relatively more collagen type 1 (i.e. collagen type 1 is more stiff than type 3 collagen) and/or more cross-linking between collagen fibrils.

In GM of spastic mice, also no differences in passive length-force characteristics were shown. For GM, passive slack length is determined by the same factors as mentioned above for SO. As physiological cross-sectional area of spastic GM was smaller, this could have contributed to a shorter GM muscle belly. Since the physiological cross-sectional area determines stiffness, the smaller physiological cross-sectional area likely caused a lower stiffness. For GM, the connective tissue content did not differ between spastic and WT mice. The observation that the passive length-force curves of spastic and WT GM were similar, is likely due to the net effects of the smaller physiological cross-sectional area on length and stiffness. The lack of difference in passive properties between spastic and WT muscles, suggest that the typical hopping gait pattern, with shorter stride lengths and less surface-paw contact is mainly caused by the overactive stretch reflex rather than by enhanced passive stiffness due to secondary morphological adaptation of triceps surea muscles.

### Effects of spasticity on muscle endurance capacity, muscle fiber type and oxidative metabolism

Spastic GM was more fatigue resistant than WT GM. This could be largely explained by the higher mitochondrial content in spastic muscles and relatively larger percentage of slow type muscle fibers. The increase in mitochondrial density and fiber type shift are in line with the observed smaller muscle fibers in spastic muscles, as the oxidative capacity of a muscle fiber is inversely related with muscle fiber size (Van Wessel et al. 2010). Chronic muscle stimulation and fatigue are physiological conditions known for their stimulatory effect on mitochondrial biosynthesis and the transition in expression of myosin heavy chain from fast to slow type (Pette and Vrbová 1999; Van Wessel et al. 2010; Hood 2001). However, based on the low physical activity of spastic mice as measured within in home-cage showed, one would not expect adaptation towards a slow and fatigue resistant muscle. A possible explanation for this phenotypical change is that spastic mouse muscle are frequently active due to the overactive stretch reflex or due to tremors.

Here we show that calf muscles of the spastic mouse show a fatigue resistant phenotype. This phenotype resembles that reported for muscles of children with spasticity, which also show higher fatigue resistance (Moreau et al. 2008). However, children with spasticity generally show lower scores on the six-minute walking test (Fitzgerald et al. 2016). This discrepancy between the muscle phenotype and endurance gait performance may be explained by impaired efficiency of the gait pattern, the limited joint range of motion, reduced balance and lower motor control in muscles from children with spasticity. Another reason may be that children with spasticity have increased motor unit firing frequencies and increased number of recruited motor units during gait (Prosser et al. 2010). This may cause fatigue and explain the discrepancy in muscle phenotype and gait performance. Whether effects of recruitment and level of muscle activation of spastic mice during gait are affected similarly as in children with spasticity remains to be determined. If any, changes in muscle phenotype with respect oxidative capacity explain the reduced gait performance (i.e. lower gait velocity) in spastic mice. However, studies on muscle fiber type distribution of human spastic muscles are not unequivocal (Dietz et al. 1986; Ito et al. 1996; Marbini et al. 2002; Rose et al. 1994; Booth, Cortina-Borja, and Theologis 2001), which is related to differences in experimental set up and type of muscles. Changes in myosin heavy chain distribution have been suggested to be dependent on the level of functional disability (Pontén, et al. 2008). Since wrist flexor muscles in children with mild spasticity were changed towards a slower phenotype, while in children with severe spasticity the muscle fiber distribution was shifted towards faster fibers (Pontén, et al. 2008). In addition, interventions aiming to reduce spasticity (e.g. Botulinum toxin injections), are also known to cause a shift myosin heavy chain distribution towards more type I muscles fibers (Dodd et al. 2005; Inagi et al. 1999; Kranjc et al. 2001). This change in myosin heavy chain distribution occurred in Botulinum toxin injected muscle as well as in the contralateral rat muscles (Dodd et al. 2005). The above illustrates the impact of severity, age and treatment and highlights that effects of spasticity are co-determined by a many other factors causing substantial variation in the characteristics of spastic human muscle.

The present results indicate that spasticity itself without any treatment causes a shift towards a slower phenotype.

### Explanations for the smaller muscle size and elevated oxidative metabolism in spastic muscle

The lower mass of spastic SO and GM were in the same order of magnitude as the percentage difference in muscle fiber cross-sectional area. This indicates that spastic muscles fibers were impaired in their radial growth. Muscle fiber growth is determined by the net rate of synthesis and degradation of contractile and structural proteins. To obtain insight in whether protein synthesis was affected by spasticity, we compared the myonuclear density and expression levels of genes involved protein synthesis and breakdown within gastrocnemius muscle, assuming that within this muscle the impact of spasticity is similar to that in SO and GM. The number of myonuclei per muscle fiber of myonuclear density was not lower in spastic muscle neither were mRNA expression levels of myostatin and insulin-like growth factor 1. These results suggest that the capacity for protein synthesis was likely not impaired at the transcriptional level. Regarding the expression of E3 ligases, there were also no differences between spastic and WT muscles. Together these data suggest that the smaller mass and physiological cross-sectional area of spastic muscles is likely the result of the relatively lower physical activity of the mice and of increased sheltering behavior causing a lower mechanical loading of the muscles. It is well known that mechanical loading stimulates the enzymatic activation of mTOR which enhances the rate of translation and protein synthesis (Van Wessel et al. 2010; Goodman et al. 2011; Hong et al. 2013; Gehlert et al. 2015). Our results suggest that during development the reduced activity of spastic mice results in a diminished mechanical loading of their muscles which is critical for their radial growth (i.e. addition of sarcomeres in parallel). The increased mitochondrial content in spastic muscles is likely explained by enhanced transcriptional activity of genes coding for mitochondrial proteins (Van Wessel et al. 2010). The observation that the number of satellite cells was not affected by spasticity indicates that spastic muscles have a typical potential for regeneration in case of injury and also that spastic muscle has sufficient potential for accretion of myonuclei required for longitudinal muscle fiber growth (White et al. 2010).

To our knowledge, muscle gene expression in children has not been extensively studied, neither mitochondrial or satellite cell content. Gastrocnemius muscle fibers of spastic children show a high variability in cross-sectional area, rather than being smaller (Ito et al. 1996), particularly in type 1 muscle fibers. In spastic children, the lack of longitudinal gracilis muscle fiber growth has been suggested to be associated with a lack of addition of sarcomeres in series as well as by myonuclear addition and loss of satellite cell numbers (Dayanidhi et al. 2015). Our results suggest that spasticity, whether or not in combination with reduced physical activity, does not alter the satellite cell population. However, it is still unknown if and how muscle extensibility plays a role in the alteration of the satellite cell population. The latter might be important, since the investigated human muscles, were undergoing lengthening surgery, because of reduced knee joint range of motion (Dayanidhi et al. 2015).

### Study limitations

Some limitations of this study should be noted. The uneven male-female ratio in both experimental groups may have increased the variation in results of the muscle fiber type distribution which could have reduced the power to confirm differences between GM of both groups. Nevertheless, endurance capacity of GM muscles was consistently higher in spastic than in WT mice, which supports a shift towards a slower muscle fiber phenotype in spastic mice which became more pronounced with age (Fig. 5).

Moreover, regarding the gene expression data, we determined target mRNA levels relative to those of the housekeeping gene. This approach does not take into account any difference in total RNA concentration within spastic and WT muscles. As high oxidative muscle fibers usually have higher RNA concentration (Van Wessel et al 2010; Habets et al. 1999) and spastic mice seem to show a more slow phenotype with smaller muscle fibers, E3 ligase and mitochondrial enzyme activity could be higher in spastic muscle and explain the observed differences in muscle size and metabolism.

## Conclusions

This study shows that glycine receptor subunit-ß deficiency in mice results in a series of motor abnormalities, including abnormal gait characterized by toe walking and hopping, impaired hind limb motor function as well as reduced physical activity. This disturbed motor function is likely the result of the overactive stretch-reflex and secondary impairment of skeletal muscle growth. Spastic mice show smaller and weaker plantar flexor muscles which becomes more apparent with age. Muscle length of GM muscle is slightly reduced, while passive length-force properties are not affected. The affected force generating capacity is explained by the smaller muscle fiber cross-sectional area, the lower number of sarcomeres in series and shorter Achilles tendon. Due to the shortening, plantar flexor muscles exert higher forces at lower muscle lengths. In contrast to the effect of maximal force generating capacity, endurance capacity of spastic muscle is improved which seems functional and may compensates for the lower force generating capacity. The glycine receptor subunit-ß deficient spastic mouse model captures multiple key features of primary effects of spasticity on muscles function that are now accurately quantified, which allows to study interventions to improve muscle functions, morphological characteristics and phenotypes in children with spasticity.

## Materials and Methods

### Animals

Mice carrying a spontaneous mutation in the LINE-1 element insertion of the glycine receptor ß-subunit (Mülhardt et al. 1994) were obtained from a cryopreserved stock at Jackson laboratories (Stock 000066; B6.Cg-Glrb^spa^/J; Bar Harbour, ME, USA). The colony was maintained on a C57Bl/6J background (Charles River Laboratories, L’Arbresle, France; European supplier of Jackson Laboratories) in the animal facilities of the Vrije Universiteit Amsterdam (Amsterdam, The Netherlands) and was backcrossed to C57Bl/6J at least every 3rd generation. Experimental homozygous Glrb^spa^ mice and wild type littermate controls (WT mice hereafter) were obtained by breeding heterozygous carriers. Food was provided on the bedding of the cage, water bottles with long spouts were used. At the age of approximately 2 – 3 weeks ear tissue samples were taken for genotyping, and mice were weaned at the age of 3 – 4 weeks. Before the age of 4 – 5 weeks approximately 25% of homozygous mice were found dead or had disappeared from the nests. These premature deaths were associated with extremely low body weight (∼5 g), whereas surviving homozygous mice had a considerable higher body weight (>7 g). A total of 15 spastic mice (8 female / 7 male) and 15 WT mice (7 female / 8 male) entered the behavioral test experiment at 4 – 5 weeks of age, and a subset of these mice were used for in situ experiments and tissue analyses (9 WT / 10 spastic mice). Additional mice were bred for in situ experiments and tissue analyses at juvenile age (10 WT and 10 spastic). Before weaning, mice were housed in IVC cages and health status of the colony was confirmed Specific Pathogen Free following FELASA 2014 recommendations. After weaning, mice were moved to a barrier with conventional open cages under SPF conditions (following FELASA 2014 recommendations) with the exception of several pathogens that are not considered harmful for the immune competent C57BL/6J background used in this study (*Helicobacter typhlonius, Pasteurella pneumotropica, Chilomastix spp, Entamoeba muris, Mouse Norovirus*). Both during breeding and experiments, mice were maintained at 12-hour light/dark cycle with lights on at 7.00 a.m., humidity between 40 – 60% and room temperature between 19 – 22 °C. Water and food were available ad libitum. Mice were group housed in macrolon type 2 short cages on sawdust bedding with cardboard curls as nesting material, except during assessment of spontaneous behavior in the automated home-cage as describes below. Behavioral experiments were performed during the light phase.

All experiments were approved by the Dutch Central Committee on Animal Experimentation (AVD112002016772) and in strict agreement with the guidelines and regulations concerning animal welfare recommended by the Dutch law.

### Study design

One cohort of 15 WT (male n= 8 / female n= 7) and 15 spastic (male n= 8 / female n= 7) mice were subjected to a series of behavioral tests at 4, 6, 10 and 14 weeks. Body weight, neurological score and righting reflex were scored every week. A subset of this first cohort of mice were used for in situ experiments and tissue analyses and included 9 WT (male n= 5 / female n= 4) and 10 (male n= 4 /female n= 6) spastic adult mice (17.9 ± 0.3 weeks). A second cohort of mice was used for in situ experiments and tissue analyses at juvenile age and included 12 WT (male n= 6 / female n= 6) and 9 spastic (male n= 6 / female n= 3) juvenile (5.4 ± 0.2 weeks).

### Spontaneous behavior

Mice were individually housed in a PhenoTyper automated home-cages for 3 days as described before, with access to water, food, and a shelter, without human intervention (Loos et al. 2014). With respect to spontaneous behaviors in the first three days in the PhenoTyper, 6 groups of behavioral parameters were defined as described below and were analysed using AHCODA (Sylics, Amsterdam, the Netherlands) with respect to temporal aspects, as described in detail before (Loos et al. 2014). Mice that build their nest outside the shelter (which interferes with calculation of all activity measures) were excluded. Mice that spend little time in the shelter (<60% of time in the shelter during light phase of day 2 and 3) in combination with being highly inactive outside the shelter (cumulative movement less than 2 cm per 5 min for >25% of time outside during light phase of day 2 and 3) were classified as sleeping outside the shelter and excluded from the analyses.

### Neurological Score and righting reflex

General wellbeing of the mice was assessed by observation of several characteristics. The neurological score was adapted from a scoring system originally developed to score the progression of mouse model of amyotrophic lateral sclerosis (Hatzipetros et al. 2015). For the neurological score, 3 items were scored; extension reflex, curling toes and tremor. With respect to extension reflex, a score of 0 was written down if a full extension of hind legs away from lateral midline was observed when mouse was suspended by its tail, and mouse could hold this for 2 s. If mice showed a partial extension reflex (score 0.5) or no extension reflex this is taken as index of neurological symptoms. With respect to tremor, trembling of hind legs during tail suspension was scored (score 0 = no tremor, score 1 = tremor present). With respect to curling toes, the position of the toes was observed during forward motion (score 0 = no curling toes, score 1 = curling toes present). To calculate the neurological score, the average score on these three items was calculated. In addition, the original scoring system described a righting test (Hatzipetros et al. 2015). When a mouse is placed on its side and is unable to right itself within 30 s from either side this is considered a high neurological score. We noticed that spastic mice were unable to right themselves occasionally, and that this could last for more than 30 s, whereas on a next occasion the same mouse would right itself quickly. Hence, the righting test score was used a separate measure, and the actual righting time was reported as result.

### Grip strength measurements

Neuromuscular function was assessed by measuring the peak amount of force (N) mice applied in grasping a pull bar connected to a force transducer (1027DSM Grip Strength Meter, Columbus instruments, Columbus, OH, USA). Mice were allowed to grasp the pull bar 5 times with front paws only, followed by grasping 5 times with front and hind paws

### Motor coordination

Motor function was evaluated using an accelerating rotarod (Roto-rod series 8, IITC Life Science, Woodland Hills, CA, USA). In the first week of testing, mice received two habituation trials of 120 s (acceleration of from 0 to 20 rpm in 120 s) followed by 3 test trials (acceleration of from 0 to 40 rpm in 180 s) on the first day, and 5 test trials on the second day. During subsequent testing weeks mice received 1 day of test trials only with 5 trials (acceleration of from 0 to 40 rpm in 180 s).

### Gait analysis

Gait measurements were recorded and analysed using the CatWalk system (Noldus IT, Wageningen, the Netherlands). Mice were introduced to the CatWalk system on day 1 for habituation, and on the second day mice were tested. Several readouts were taken. Maximum Contact Area, which is the maximum area of a paw that comes into contact with the glass plate. Intensity ranges from 0 to 255. The intensity of a print depends on the degree of contact between a paw and the glass plate and increases with increasing weight. Therefore, intensity is a measure of weight put on the glass plate. Print length, which is the length (horizontal direction) of the complete print. The complete print is the sum of all contacts with the glass plate, as if the animal’s paw would have been inked. Print width, which is the width (vertical direction) of the complete paw print. Print area, which is the surface area of the complete print. Stride length, which is the distance between successive placements of the same paw.

The Regularity Index, which expresses as the number of normal step sequence patterns relative to the total number of paw placements. Base of Support, which is the mean distance between either front paws or hind paws. There were several exclusion criteria: Non-compliant runs as defined by the CatWalk software (Minimum run duration: 0.5 seconds, Maximum run duration: 5.0 seconds, Minimum number of compliant runs: 3 runs, Minimum number of consecutive steps: 6, Average speed between: 10 and 60 cm/s)

### Surgery and preparation for in situ muscle function measurements

Mice received a subcutaneous injection 0.1 mg·kg^−1^ Temgesic^®^ (Buprenorphine, Reckeitt Benckiser, UK) as an analgesic 20 minutes prior to surgery. Subsequently, they were anaesthetized with 4% Isoflurane, 0.2 L·min^−1^ O2 and 0.2 L·min^−1^ air. After nociceptive reflexes ceased, Isoflurane levels were maintained at 0.5-2.0%. Body temperature was maintained at 35 °C using an electrical heating pad. To prevent the exposed muscles from dehydration, the lower leg was placed in a self-irrigating humidifier with a constant temperature of 35 °C and a humidity of 80-90%. On regular basis the exposed tissue was observed and when needed irrigated with isotonic saline.

SO and GM were dissected free from surrounding tissue and from distal insertion while blood supply and innervation were maintained. Cuff electrodes were placed around GM and gastrocnemius lateralis branches of the TA nerve, and served to stimulate the muscles. Finally, the femur was fixated at the distal condyle by a clamp. Fig. 2 shows an overview of the experimental setup.

### Experimental setup and in situ muscle force measurements

To attain a full characterization of the calf muscles, we assessed for SO and GM the following variables: a) length-force relationship, b) frequency-force relationship, c) fatigability, and d) rate of maximal force development. Contractions for all protocols were induced by supramaximal electrical stimulation of GM and gastrocnemius lateralis nerves for either SO or GM muscle, respectively. For assessment of the length-force relationship tetanic contractions (150 Hz, 300 ms) were evoked at increasing muscle lengths (0.5 mm stepwise increase). Each tetanus was preceded by two twitches, to allow adjustment of muscle length. After each contraction muscles were allowed to rest for 2 minutes. Muscles were lengthened to about 1 mm over their optimum length. Muscle/muscle-tendon complex length were calculated by measuring the distance between the markers placed on the proximal and distal parts of the muscle on the images made with a Panasonic HC-V720 camera. Two minutes after finishing the length-force protocol muscles were stimulated by applying a 400 Hz, 150 ms pulse train to determine the rate of maximal force development. The frequency-force protocol was started after a recovery period of two minutes at a short length. The muscle was stimulated at optimum length with increasing frequencies starting at 5 Hz and ending at 250 Hz each time during 300 ms. Finally, SO and GM muscles were stimulated once every second for 2 minutes to test their fatigue resistance (SO: 330 ms pulse train at 100 Hz, GM: 150 ms pulse train at 30-80 Hz, a frequency corresponding with 40% of optimum force). The mouse was sacrificed by an overdose of 20% Euthasol^®^ 0.2ml injected intracardially. Plantar flexor muscles together with TA were carefully removed and aligned according to their muscle ?bre arrangement (Jaspers et al., 1999). The right-sided SO and GM muscles were covered with a thin layer of Tissue-Tek (Jung, Leica, Mycrosystems, Germany) and frozen in liquid nitrogen. The right gastrocnemius lateralis was frozen for gene-expression analysis. The left sided plantar flexor muscles and tibialis anterior were fixed and stored.

### Morphometric analyses

Serial number of sarcomeres within muscle fibers was determined in left SO and GM. At least 4 proximal and 4 distal muscle fibers were isolated and mounted on glass slides with 50% glycerol, covered with a coverslip and sealed with nail polish as described by (Heslinga and Huijing 1990). Images were taken of every single fiber using a bright field microscope and a 10x objective (Axioskop 50 microscope, Zeiss). Mean sarcomere length was determined by the number of sarcomeres over the entire muscle fiber length. The number of sarcomeres per fiber was calculated by dividing the measured muscle fiber length by the mean sarcomere length of the muscle fiber. Optimum muscle fiber length was calculated conform Equation 1, optimum sarcomere length (2.2 µm), was taken as the mean value of the optimum sarcomere length in the length-force curve as described in (Edman 2005) and multiplied by the number of sarcomeres in series. Physiological cross-sectional area (PCSA) was calculated conform Equation 2, where muscle volume was obtained by Equation 3. To obtain muscle volume the muscle weight was multiplied by muscle density (ρ), which was set on 1.0597 g·cm^−3^, as suggested in Mendez and Keys (1960).

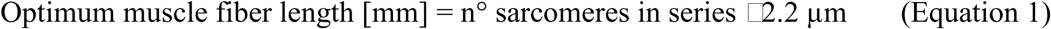

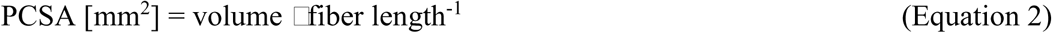

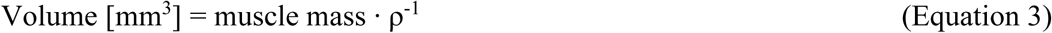

The tendon length at optimum muscle length was obtained from the images taken during length-force measurements by subtracting muscle belly length from GM muscle-tendon complex length at optimum length.

### Cryosectioning and Histology

Serial cross-sections (10 µm) were cut from the mid-belly of SO and GM using a cryostat (MICROM HM 550) at −20 °C. Sections were mounted on glass slides (Menzel-Gläser, superfrost^®^ plus, Germany), air dried and stored at −80 °C until further use. Three cross-sections were stained for succinate dehydrogenase (SDH) after 15 minutes of air drying, as described previously (Van der Laarse, et al. 1989). SDH activity was expressed as the absorbance at 660 nm·μm^−1^, section thickness per second of incubation time (ΔA660·μm^−1^·s^−1^).

### Immunohistochemical staining of myosin heavy chain isoform expression

Serial sections of SO and GM were immunohistochemically stained against type I, IIA, IIB, and IIX Myosin heavy chain (MHC) using primary monoclonal antibodies BAD5 (1 μg·Ml^−1^), SC-71 (1 μg·Ml^−1^), 6H1 (10 μg·Ml^−1^), and BF-F3 (1 μg·Ml^−1^) (Developmental Studies Hybridoma Bank, USA), respectively. After air drying for 10 minutes, sections were blocked with 10% Normal Goat serum for 60 minutes. Sections were incubated with a cocktail of two primary antibodies for 60 min. Subsequently, sections were washed in PBS three times for 5 min and incubated in the dark with a cocktail of two secondary antibodies Alexa Fluor IgG2b 647/IgM 488, IgG1 488/IgM647 (Life Technologies, The Netherlands) for 60 min. After washing with PBS 3 times 5 min in the dark, sections were incubated with wheat germ agglutinin (WGA, 1:50, Life Technologies, The Netherlands) for 20 min, and washed once more with PBS 3 times 5 min. Finally, Sections were enclosed with Vectashield^®^-hardset mounting medium with DAPI (1.5 μg·Ml^−1^, Vector Laboratories, USA). Images were captured at 10× magnification using a CCD camera (PCOl Sensicam, Kelheim, Germany) connected to a fluorescence microscope (Axiovert 200M; Zeiss, Göttingen, Germany) with image processing software Slidebook 5.0 (Intelligent Image Innovations, Denver, Colorado).

Muscle fiber cross-sectional area was determined as the mean cross-sectional area of at least 50 fibers. For SO muscle fiber cross-sectional area was determined per muscle fiber type, while for GM fiber cross-sectional area was determined in high and low oxidative regions without making a distinction between muscle fiber types.

### Quantification of intramuscular connective tissue

Intramuscular connective tissue was determined and quantified as described previously (De Bruin et al. 2014). The sections were air-dried for 10 min and fixed with Acetone for another 10 min at −20°C. Subsequently the sections were fixated with Bouin for 30 min and stained in Sirius Red saturated with picric acid for 30 min. After the sections were washed in 0.01N HCl, they were twice rinsed in ethanol absolute and twice in Xyleen (Bisolve Chimie, SARL, France). Thereafter, sections were enclosed with Entellan^®^ (Merck, the Netherlands). Binary images were taken of the stained cross-section, so that red areas in the cross-section were covered with black in the image.

The endomysium, connective tissue surrounding a single muscle fiber (MF), was quantified by selecting 10 regions containing only endomysium and muscle fibers. Absolute endomysium (A_E_) was normalized to fiber cross-sectional area (A_E_ divided by the number of MF in the selected area). The mean endomysium thickness was measured separately.

Three domains of perimysium were measured separately: length of collagen surrounding the smallest muscle fiber fascicles (L_p1_), length of larger fascicles which contain several primary fascicles (L_p2_) and length of collagen which did traverse the cross-section (L_p3_; traversing the cross-section). Thickness were manually measured every 25 µm for at least 5 fascicles in L_p1_ and L_p2_ and 25 µm over a length of 1 mm. Since many of the cross-sections were cleft, the thickness of the perimysium was not always measurable in each of the three domains.

### Myonuclear density and satellite cell number per muscle fiber

To quantify myonuclear density and muscle stem cells number per muscle fiber were counted, sections were co-stained with DAPI and anti-Pax7 antibodies according to van der Meer et al. (2011). Sections were fixed in 4% formaldehyde in PBS for 10 min, washed with PBST and blocked in 10% normal goat serum in PBS for 30 min. Subsequently, sections were incubated in 0.1% bovine serum albumin–PBS solution containing 4 μg·mL^−1^ Pax7 antibody (Developmental Studies Hybridoma Bank, USA) in the dark for 60 min, washed in PBST, incubated with Alexa Fluor 488 (1:500, Molecular Probes, Life Technologies) goat anti-mouse secondary antibody (Invitrogen, Breda, The Netherlands) and washed with PBST. Then sections were incubated in the dark for 20 min with Texas red-WGA (1:50) conjugate (Invitrogen, Breda, The Netherlands) and washed with PBS. Finally, sections were enclosed with Vectashield®-hardset mounting medium with DAPI (Vector Laboratories, USA). Images were captured at 10× magnification using a CCD camera (PCOl Sensicam, Kelheim, Germany) connected to a fluorescence microscope (Axiovert 200M; Zeiss, Göttingen, Germany) with image processing software Slidebook 5.0 (Intelligent Image Innovations, Denver, Colorado).

The number of myonuclear fragments per muscle fiber cross-section was manually counted in an average of 50 fibers per muscle, while muscle stem cell fragments per muscle fiber cross-section were counted throughout the cross-section. To calculate the number of myonuclei (Mn) in muscle fiber, we used equation 4, as suggested in (Jaspers et al. 2006).

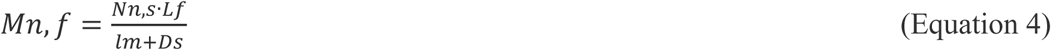

Where N_n,s_ is the counted myonuclei per fiber cross-section, L_f_ the muscle fiber length, L_m_ is the length of the myonuclei, and D_s_ the thickness of the cross-section. The length of myonuclei was assumed to be 12 µm (van der Meer et al. 2011). The myonuclear domain was obtained by dividing the fiber cross-sectional area by the myonuclei per 1 mm of fiber length. Number of muscle stem cells is reported as counted fractions per muscle fiber cross-section of SO and GM muscles.

### RNA isolation and real-time PCR

RNA was extracted from frozen gastrocnemius lateralis muscles using TRIZOL (Invitrogen, USA) and homogenized with an automatic potter homogenizer (B. BRAUN, Germany) followed by RNA purification using the Ribo pure-kit (Thermo Fisher Scientific, Waltham, MA). RNA concentration and purity were measured using spectroscopy (Nanodrop Technologies, Wilmington, DE). RNA was reversed transcribed using SuperScript(tm) VILO(tm) MasterMix (Applied Biosystem, Carlsbad, CA) and the thermal cycler 2720. For each PCR target, 12.5 ng of the RT reaction product was amplified in duplicate using Fast SYBR Green master mix (Applied Biosystems). Real-time PCR was performed on a StepOne Real-Time PCR system (Applied Biosystems). For MuRF1, MAFbx, Mstn, IGF-1 Ea, MyoD, Pax7 and SDH mRNA, cycle threshold was subtracted from the mean cycle threshold value of 18S rRNA ΔCt and converted into relative concentrations by 2^−ΔCt^. The sequences for the primers (Invitrogen, The Netherlands) used for the specific RNA and mRNA targets are shown in Table 8.

**Table 8:**
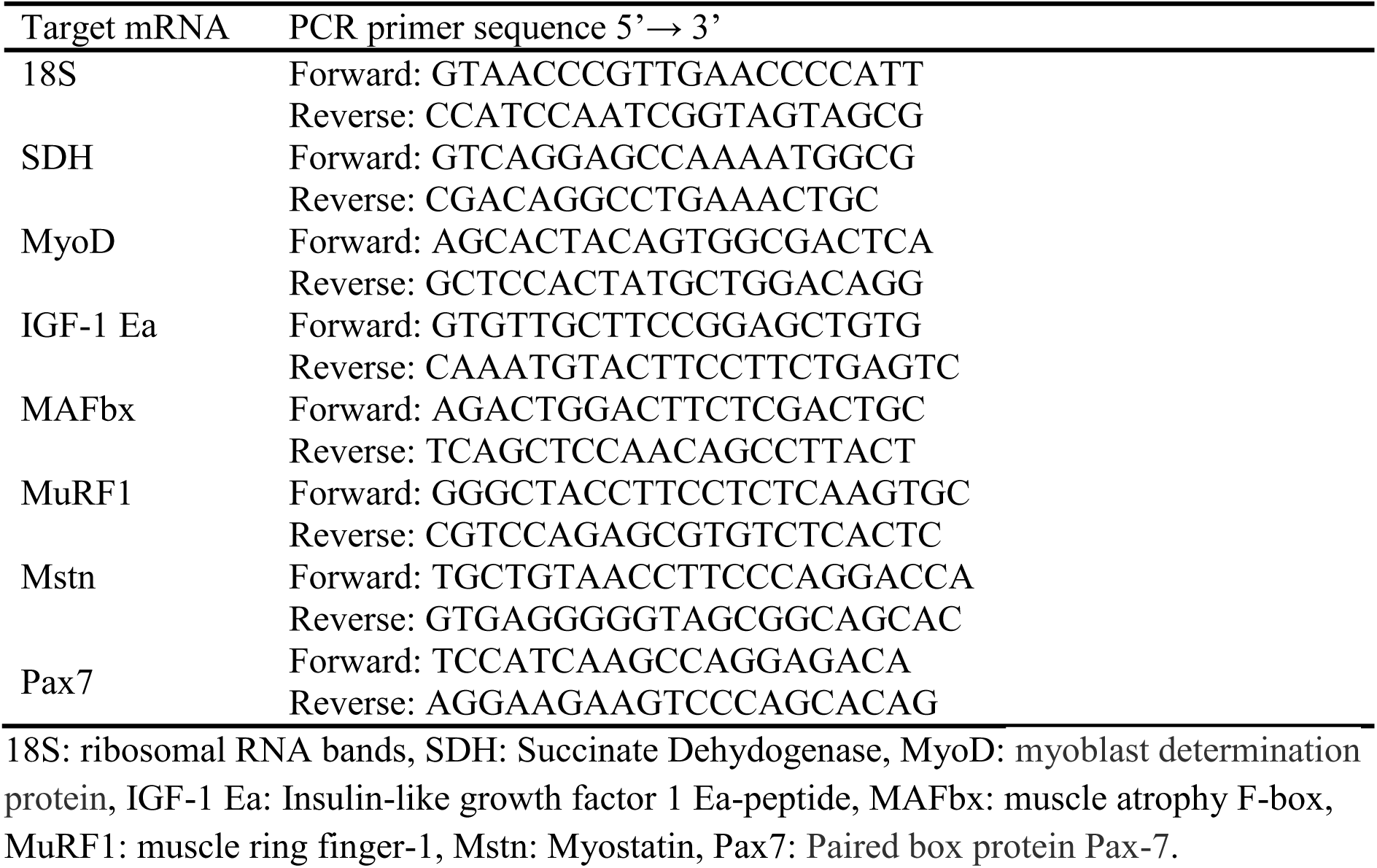
Sequence of the specific primers used in the quantitative PCR analyses

### Data treatment

#### Length-force characteristics

Passive muscle force as a function of muscle length was obtained by the mean force between the second twitch and the tetanus and to a second order exponential (Equation 5). Active muscle force was calculated by subtracting passive force at each muscle length from total muscle force. The active length-force was fitted by a polynomial function (Equation. 6).

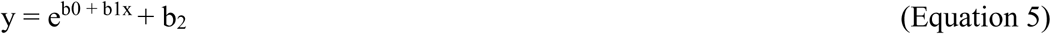

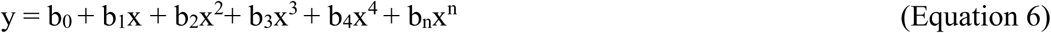

In equations 5 and 6, b_0_ to b_n_ represent constants determined by the least square fitting procedure. To select the polynomial order that describes the length-force relationship most adequately, fits between a 2^nd^ and a 5^th^ order were tested using one-way ANOVA. The lowest polynomial order that yielded a significant improvement of the LF fit was elected. Muscle optimum length and muscle active slack were defined as the muscle lengths at which the maximum of the polynomial was encountered, and the intercept of the fitted curve with the x-axis, respectively.

#### Force-frequency relationship

Calcium sensitivity in SO and GM were assessed by electrical muscle stimulation over a range of frequencies (i.e. 5 to 250 Hz in adult and 10 to 125 Hz in juvenile mice) at length of the active muscle at optimum length. Force-frequency relationship was obtained by subtracting the mean passive muscle force (measured between the second twitch and the tetanus) from the total force and dividing total force at each frequency by the maximal force.

#### Excitation-contraction coupling

Maximal rate of force development was described as the rate at which the active force went from zero to maximal active force and is expressed in Nm·ms^−1^.

#### Endurance capacity

Fatigability was assessed by calculating the decrease in active muscle force at the end of the series of contractions and express this as percentage of the maximal obtained force during the protocol.

Matlab scripts were used for evaluation of the force data, and for calculation of number of sarcomeres in series (version R2017a, The MathWorks, Inc., Massachusetts, USA). The public domain software ImageJ (version 1.52e, US National Institutes of Health, Maryland, USA) was used to assess the quantity of endomysium, perimysium, number of myofiber, type, fiber CSA and mean sarcomere lengths.

#### Statistics

Data were tested for normality and homogeneity and in case assumptions were not met, data was transformed (i.e. (natural)-log, exponential, or square-root transformation).

In juvenile mice there was a significant difference in age between males and females, which was reflected in body and muscle masses. Therefore, we performed the statistics for all variables except force-frequency, sarcomeres in series and histological parameters only on juvenile male mice.

Two-way mixed ANOVA was used to test physical performance, gait, length-force and force-frequency data. In case effects of gender could not be tested, independent t-tests between group (juvenile mice). Gender was also excluded when the amount of pairwise comparisons excluded the actual number of mice per group (e.g. in fiber types of GM muscle). Interaction effects and gender main effects were explicitly described when present, otherwise only group main effects were reported. General estimating equations was used to determine the relationship of righting time and Neurological Score and righting reflex with age and age*gender within the spastic mice group. These statistics did not include WT mice, as their score was constantly zero. Two-way ANOVA were used to calculate all the other parameters. All statistical analyses were performed using SPSS (IBM SPSS statistics 25, New York, USA). Prism (version 8.2.0, GraphPad software Inc, California, USA) was used in order to plot the figures. Effects were considered significant at p<0.05. Data is presented as mean ± SEM.

Behavioural data of this study will be published in the public repository of AHCODA-DB at https://public.sylics.com.

## Supplementary data

**Supplementary figure 1.**
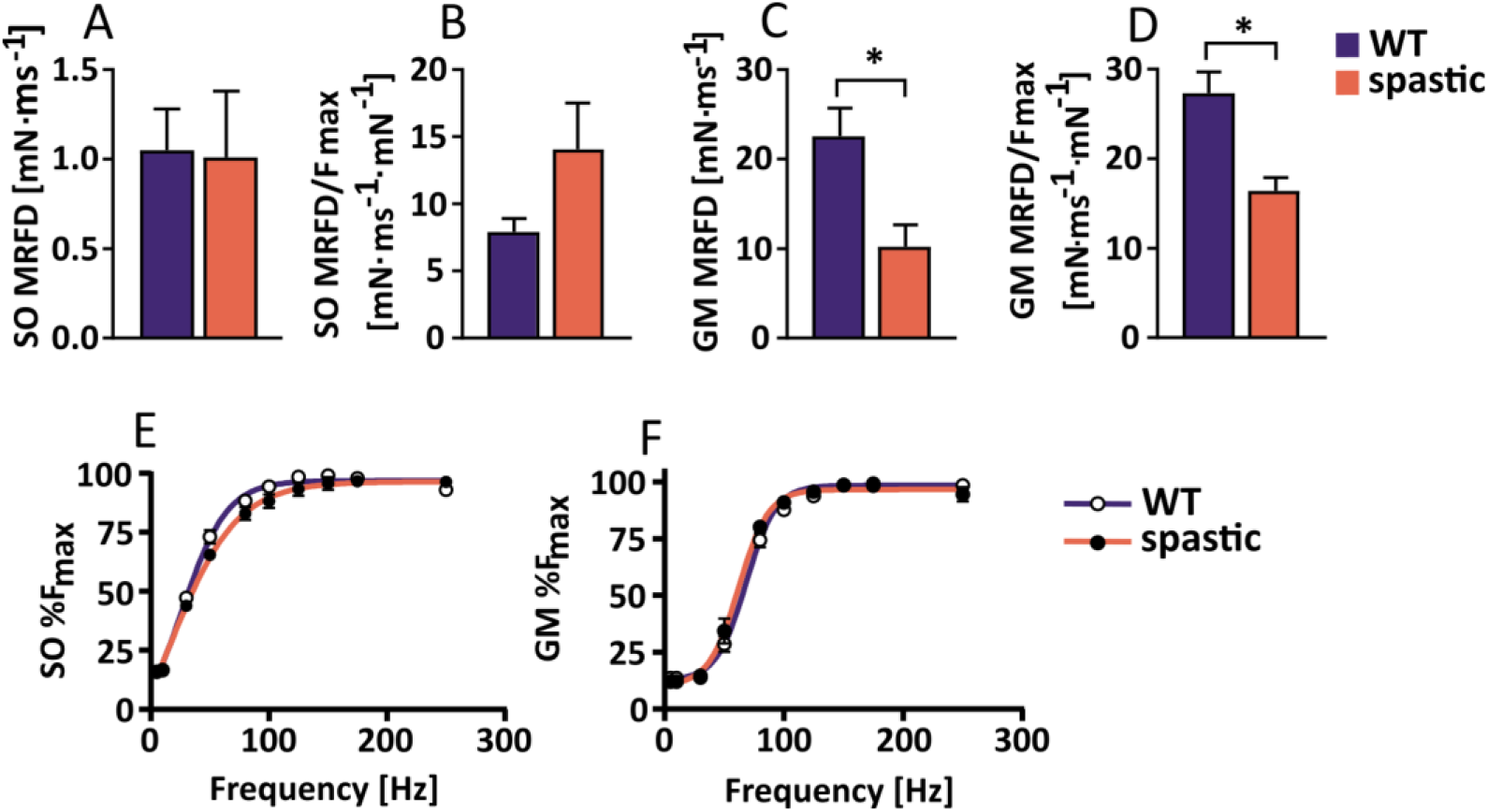
Contractile properties of juvenile spastic SO and GM muscles. (A) Absolute maximal rate of force development of SO muscle, (B) Rate of maximal force development of SO normalized for maximal force, (C) Absolute rate of maximal force development of GM muscle, (D) Rate of maximal force development of GM, (E) Force-frequency relationship of SO muscle, (F) Force-frequency relationship of GM muscle.

**Supplementary figure 2.**
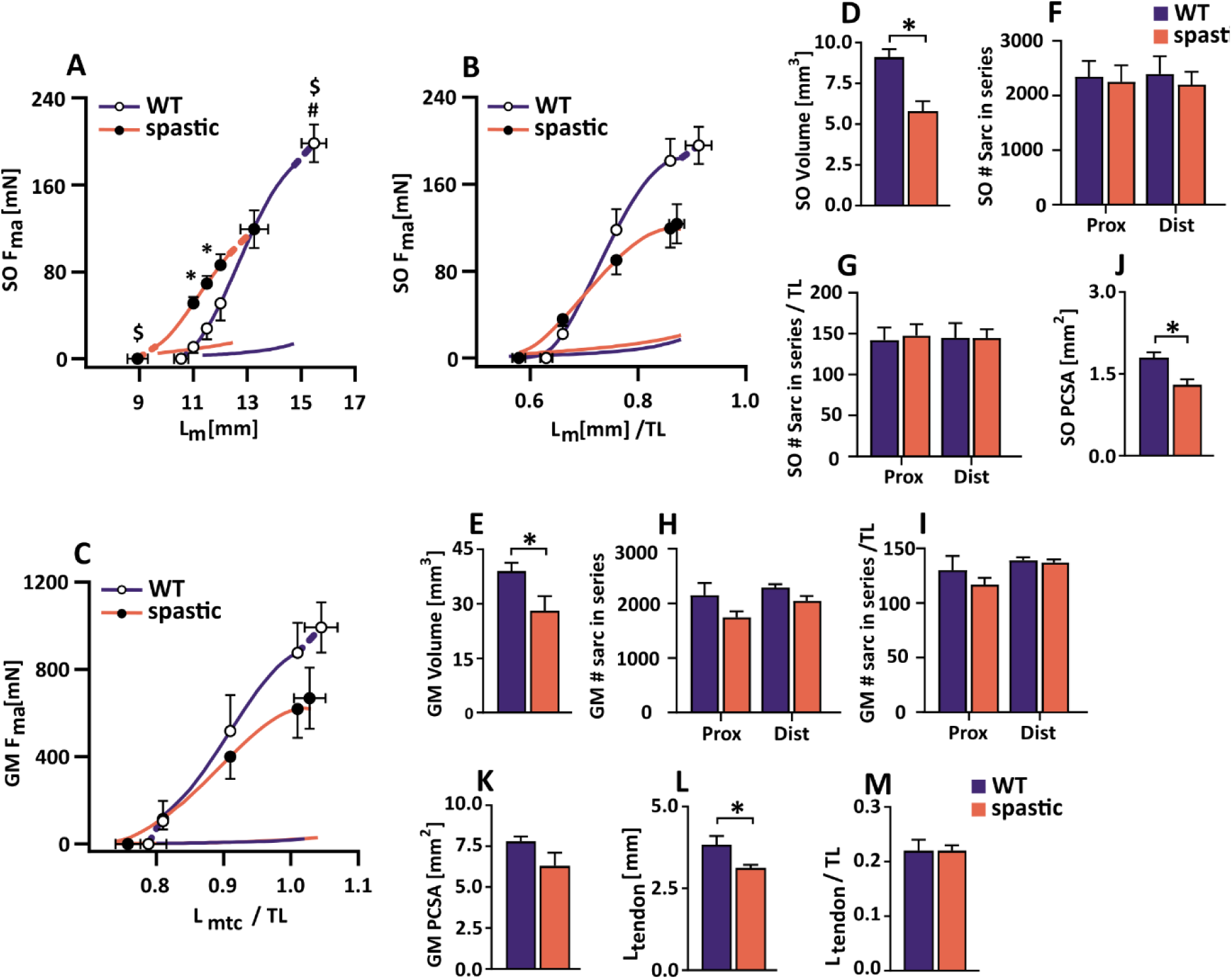
Effects of spasticity on juvenile muscle length-force relationship as well as determinants of muscle length and muscle force. (A) SO absolute length-force, (B) SO length-force normalized for tibia length, (C) GM length-force normalized for tibia length, SO muscle volume, (E) GM muscle volume, (F) SO number of sarcomeres in series of proximal (Prox) and distal (Dist) fibers, (G) SO number of sarcomeres in series of Prox and Dist fibers per unit of tibia length, (H) GM number of sarcomeres in series of Prox and distal Dist fibers, (I) GM number of sarcomeres in series of Prox and Dist fibers per unit of tibia length, (J) SO PCSA, (K) GM PCSA, (L) Achilles tendon length, (M) Achilles tendon length normalized for tibia length.

**Supplementary figure 3.**
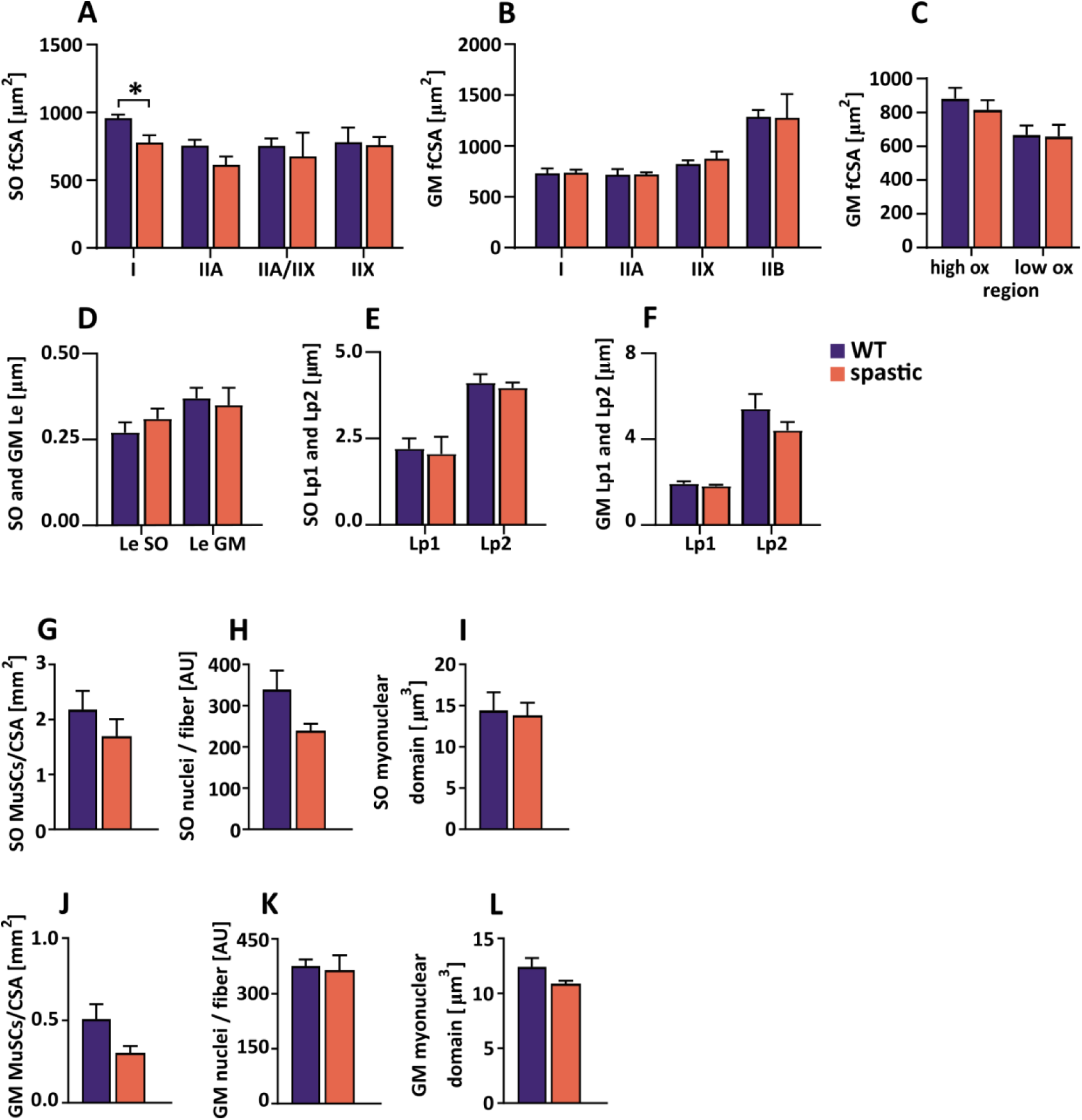
Active and passive muscle force determinants of adult spastic muscles. (A) SO fiber cross-sectional area per fiber type, (B) GM fiber cross-sectional area per fiber type, (C) GM fiber CSA per muscle region, (D) SO and GM endomysium length, SO perimysium length, (F) GM perimysium length, (G) SO GM muscle stem cells per cross-sectional area, (H) SO nuclei per fiber, (I) SO myonuclear domain, (J) GM muscle stem cells per cross-sectional area, (K) GM SO nuclei per fiber, (L) GM myonuclear domain.

**Supplementary figure 4.**
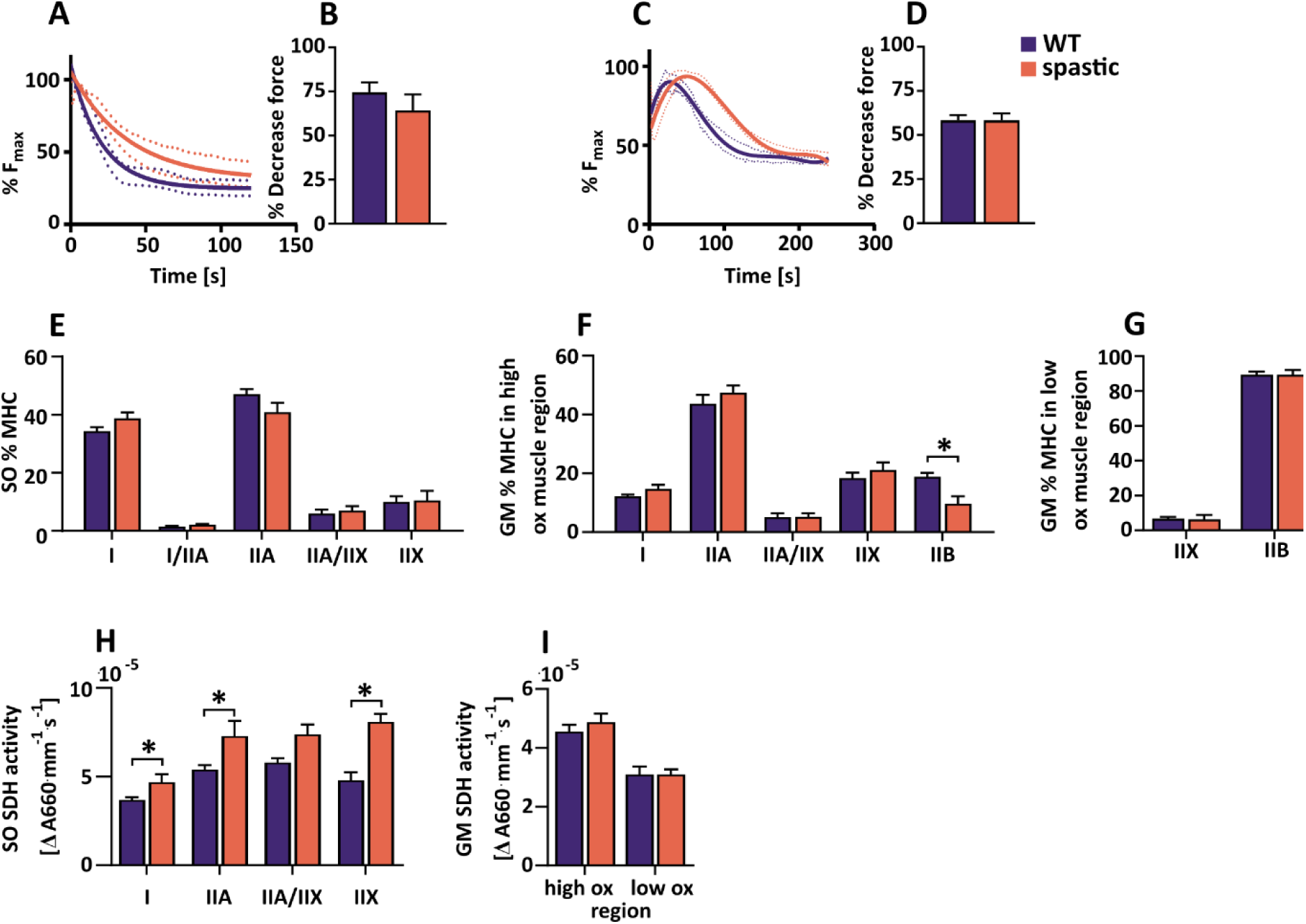
Endurance capacity of juvenile SO and GM as well as endurance determinants. (A) SO endurance curve, (B) SO percentage of decreased force at the end of the fatigue protocol, (C) GM endurance curve, (D) GM percentage of decreased force at the end of the fatigue protocol, (E) SO MHC distribution, (F) GM MHC distribution in the low oxidative muscle region, (G) GM MHC distribution in the high oxidative muscle region, (H) SO SDH activity, (I) GM SDH activity.

